# Evidence for influenza B virus hemagglutinin adaptation to the human host: high cleavability, acid-stability and preference for cool temperature

**DOI:** 10.1101/736678

**Authors:** Manon Laporte, Annelies Stevaert, Valerie Raeymaekers, Talitha Boogaerts, Inga Nehlmeier, Winston Chiu, Mohammed Benkheil, Bart Vanaudenaerde, Stefan Pöhlmann, Lieve Naesens

## Abstract

Influenza A virus (IAV) and influenza B virus (IBV) cause yearly epidemics with significant morbidity and mortality. When zoonotic IAVs enter the human population, the viral hemagglutinin (HA) requires adaptation to achieve sustained virus transmission. In contrast, IBV has been circulating in humans, its only host, for a long period of time. Whether this entailed adaptation of IBV HA to the human airways is unknown. To address this question, we compared seasonal IAV (A/H1N1 and A/H3N2) and IBV viruses (B/Victoria and B/Yamagata lineage) with regard to host-dependent activity of HA as the mediator of membrane fusion during viral entry. We first investigated proteolytic activation of HA, by covering all type II transmembrane serine protease (TTSP) and kallikrein enzymes, many of which proved present in human respiratory epithelium. Compared to IAV, the IBV HA0 precursor is cleaved by a broader panel of TTSPs and activated with much higher efficiency. Accordingly, knockdown of a single protease, TMPRSS2, was sufficient to abrogate spread of IAV but not IBV in human respiratory epithelial cells. Second, the HA fusion pH proved similar for IBV and human-adapted IAVs (one exception being HA of 1918 IAV). Third, IBV HA exhibited higher expression at 33°C, a temperature required for membrane fusion by B/Victoria HA. This indicates pronounced adaptation of IBV HA to the mildly acidic pH and cooler temperature of human upper airways. These distinct and intrinsic features of IBV HA are compatible with extensive host-adaptation during prolonged circulation of this respiratory virus in the human population.

**Importance:** Influenza epidemics are caused by influenza A (IAV) and influenza B (IBV) viruses. IBV causes substantial disease, however it is far less studied than IAV. While IAV originates from animal reservoirs, IBV circulates in humans only. Virus spread requires that the viral hemagglutinin (HA) is active and sufficiently stable in human airways. We here resolve how these mechanisms differ between IBV and IAV. Whereas human IAVs rely on one particular protease for HA activation, this is not the case for IBV. Superior activation of IBV by several proteases should enhance shedding of infectious particles. IBV HA exhibits acid-stability and a preference for 33°C, indicating pronounced adaptation to the human upper airways, where the pH is mildly acidic and a cooler temperature exists. These adaptive features are rationalized by the long existence of IBV in humans, and may have broader relevance for understanding the biology and evolution of respiratory viruses.

## Introduction

The global burden of seasonal influenza is estimated at 3 to 5 million severe cases (1) and 290,000-650,000 fatalities per year (2). In addition, influenza A virus (IAV) causes sporadic pandemics with an even more severe impact on public health. Influenza B virus (IBV) does not cause pandemics but is responsible for a significant proportion of seasonal influenza (3). IBV may account for 22-44% of influenza deaths in children (3) and also elderly persons are at high risk (4). Between 1997 and 2009, IBV was the predominant cause of influenza-associated death in 4 out of 12 seasons (5). It was estimated that IAV and IBV diverged approximately 4,000 years ago (6). Current IBV strains fall into two lineages, B/Victoria and B/Yamagata, which diverged about 40 years ago and are now cocirculating globally (7). Gene reassortment between the two lineages is common yet not seen for the HA, PB1 and PB2 gene segments (7). Whereas IAV is widespread in birds and mammals (8), IBV is restricted to humans with no sustained animal reservoir (9) and only rare spillover events (10). When a zoonotic IAV enters the human population, adaptive changes in multiple viral proteins are required to achieve sustained human-to-human transmissibility (11). Which adaptive features were acquired by IBV as a result of its long existence in humans has hardly been investigated (9).

One major protein involved in this host adaptation is the viral hemagglutinin (HA), the mediator of viral entry (12, 13). HA binds to sialylated glycans on the cell surface to enable virus uptake by endocytosis. Next, HA mediates fusion of the viral envelope and endosomal membrane, allowing release of the viral genome into the cytoplasm. Membrane fusion occurs when virus-containing endosomes mature from early (pH ∼6) to late (pH ∼5) endosomes, providing the low pH trigger for drastic refolding of HA and expulsion of its fusion peptide (14). To gain membrane fusion-competence, HA relies on host cell proteases for posttranslational cleavage of the HA0 precursor into two polypeptides, HA1 and HA2, a process termed priming or activation (15). In human IAV and IBV strains, HA0 has a monobasic (single arginine) cleavage site that is recognized by trypsin-like serine proteases. Understanding the protease dependency of HA might reveal biological differences between IAV and IBV, and direct drug concepts to target these proteases for influenza therapy (16, 17). A series of proteases have been proposed for IAV (reviewed by Böttcher-Friebertshäuser *et al*. (18)), but for IBV hardly any information is available. The type II transmembrane serine protease (TTSP) TMPRSS2 (transmembrane protease serine 2) is an activator of IAV HA0 in cell culture and essential for IAV replication in mice, at least for some IAV subtypes (19–22). In humans, a gene polymorphism leading to higher *TMPRSS2* expression was correlated with the risk for developing severe influenza (23). In cell culture, IAV can also be activated by some other TTSPs and kallikreins, but which of these support spread of IAV in human airways remains to be demonstrated [reviewed in: (16, 24)]. Regarding IBV, TMPRSS2 was shown to cleave IBV HA0 (17) and mediate virus spread in some human airway cell culture models (25), however this protease appeared dispensable for IBV pathogenesis in mice (26). Hence, the protease recognition profile of IBV HA is largely unknown.

Two other adaptive features of HA are related to its pH- and temperature-dependence. Again, data for IBV are very limited. Successful virus replication and transmission require a balance between the low pH that triggers HA refolding and membrane fusion during viral entry, and acid-stability of progeny virus in the respiratory tract and environment (27, 28). Extracellular virions are sensitive to inactivation in mildly acidic parts of the upper respiratory tract (URT) like the nasal cavity (27, 29, 30). The acid-stability of HA appears to increase when a zoonotic IAV enters the human population and evolves into a human-to-human transmissible strain (31, 32). Studies in ferrets showed that increased IAV HA acid-stability contributes to airborne transmissibility (31, 33, 34). In addition, human airways exhibit a temperature gradient, from ∼30-32°C in the nasal mucosa (35), to ∼32°C in the upper trachea and ∼36°C in the bronchi (36). Avian IAVs, adapted to the temperature of the avian enteric tract (40°C), show restricted growth at cooler temperatures (∼32°C). Whereas the temperature dependency is fairly understood for the PB2 subunit of the viral polymerase (37), this is not the case for the viral surface glycoproteins, HA and neuraminidase (38). It is conceivable that HA proteins of IBV or human IAV might show intrinsic adaptation to the temperature of human airways, including the cooler temperature of the URT.

To understand how these human airway-specific factors may influence the membrane fusion activity of HA, we here compared the HA proteins of four seasonal human IAV and IBV viruses. Their proteolytic activation was analyzed by covering all members of the human TTSP (39) and kallikrein (KLK) families (40). We demonstrate that IBV HA exhibits much more efficient activation by a broad range of airway proteases, explaining why TMPRSS2 alone was required for spread of IAV but not IBV in human respiratory epithelial cells. Whereas the HA proteins of IBV and human IAV showed similar low pH-dependence to trigger membrane fusion, a distinction was seen in terms of temperature dependence, with IBV HA having clear preference for 33°C. For one of the two IBV lineages, this cooler temperature was required to achieve membrane fusion. We propose that these distinct and intrinsic properties of the IBV HA protein might reflect extensive adaptation of IBV to the human respiratory tract.

## Results

### Human lung tissue and airway epithelial cell lines are rich in TTSP and KLK enzymes

Before investigating which TTSPs or KLKs are involved in HA activation of IAV or IBV, we assessed expression of these proteases in human airways. Their mRNA levels were quantified in tissue samples of healthy human lungs (from eight different donors; Figs 1A and 1B) and three continuous cell lines (Calu-3, 16HBE and A549) derived from human airways (Figs 1C and 1D). Lung tissue was shown to express abundant levels of *TMPRSS2* and *ST14/matriptase*. All other TTSPs showed high and comparable transcript levels, except for *TMPRSS11F* and *TMPRSS15* which were present at lower levels (Fig. 1A: left). These results agreed with microarray data for 108 healthy human lungs, retrieved from the GEO database (Fig. 1A: right). Human lung tissue contained high mRNA levels for several KLKs, but these showed more variable expression compared to the TTSPs (compare right panels of Figs 1A and 1B). Calu-3 and 16HBE cells (Figs 1C and 1D) showed abundant expression of *ST14/matriptase*, and Calu-3 cells also contained high levels of *TMPRSS2, TMPRSS3* and *TMPRSS4.* Both cell lines expressed several *KLK* transcripts. In A549 and HEK293T cells, expression of these proteases was generally lower. As expected, human lung tissue and the four cell lines abundantly expressed *FURIN*, which recognizes multibasic HA0 cleavage sites of highly pathogenic avian IAVs (41) (data not shown).

**Figure 1.**
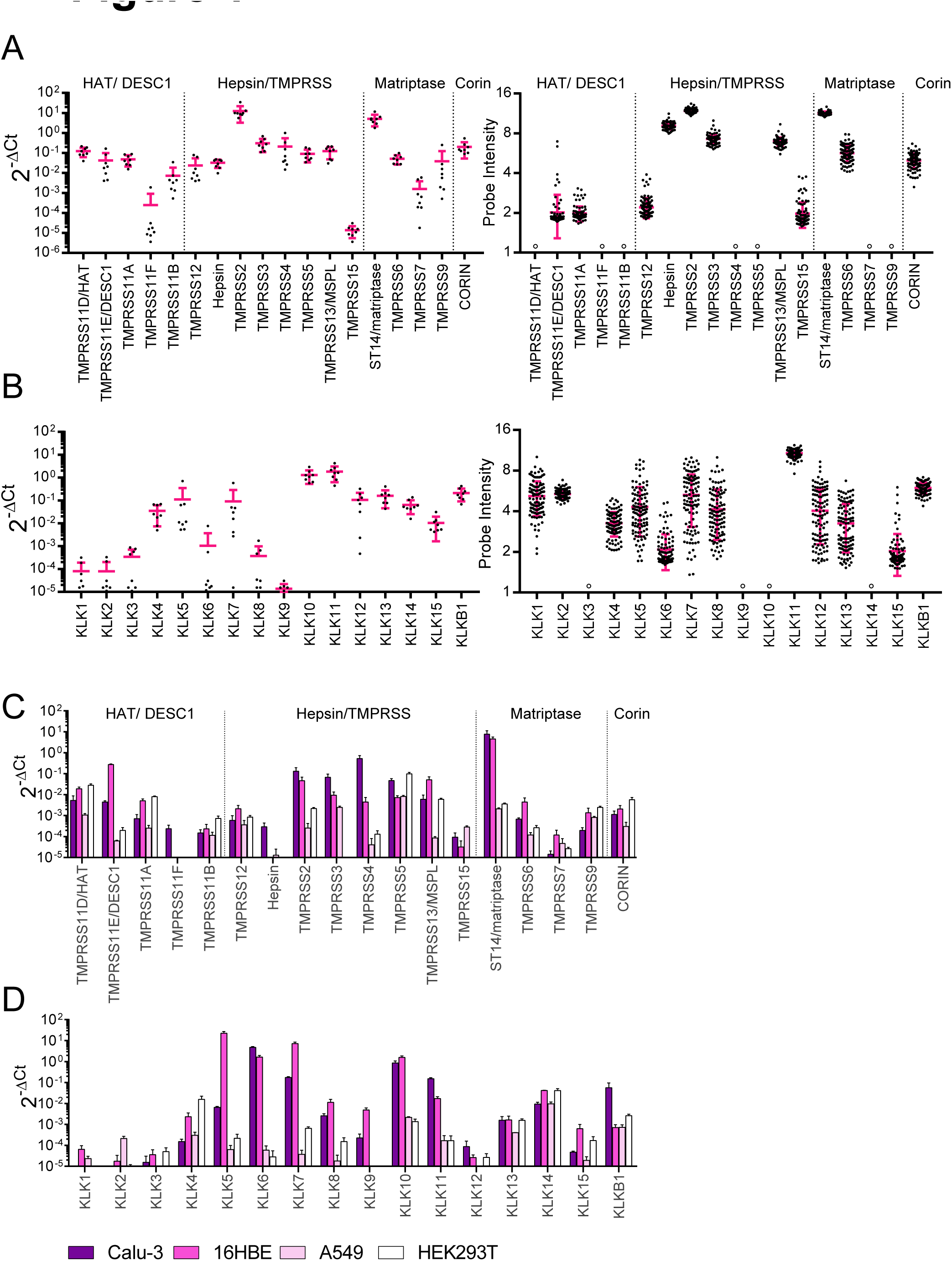
Human lung tissue and airway epithelial cell lines are rich in TTSP and KLK enzymes. (A) *TTSP* and (B) *KLK* transcript levels in healthy human lung, measured by RT-qPCR on samples from eight different donors (left), or retrieved from the NCBI GEO database (GEO dataset GSE47460) (right). Individual data points (in black) ± SEM (in pink). °No data available. (C, D) Transcript levels in human airway epithelial Calu-3, 16HBE and A549 cells, and human embryonic kidney HEK293T cells. The four TTSP subfamilies are indicated at the top of the graphs.

### IBV HA0 is efficiently cleaved by a broad range of TTSPs

To investigate which of these proteases can activate the HA proteins of IAV or IBV, expression plasmids were generated for all 18 human TTSPs and 16 KLKs, bearing a C-terminal flag tag (for details see Table S1). When we analyzed expression in transfected cells, most proteases showed a protein level similar to that of TMPRSS2 regarded as the reference (Fig. S1C). For TMPRSS11B, TMPRSS12, TMPRSS7 and KLK13, expression was three-fold lower. For TMPRSS11D/HAT and TMPRSS11E/DESC1 expression was very weak, meaning that the results from the HA cleavage assay are an underestimate for these two proteases. In this assay, the proteases were co-expressed with the HAs from four seasonal IAV (A/H1 and A/H3) and IBV (B/Yam and B/Vic) viruses (Fig. 2A). HA0 cleavage was assessed by western blot for the HA1 or HA2 cleavage product, depending on which epitope was recognized by the anti-HA antibody. The % cleaved HA was calculated relative to the trypsin control, consisting of cells transfected with HA plus empty plasmid, and briefly exposed to exogenous trypsin.

**Figure 2.**
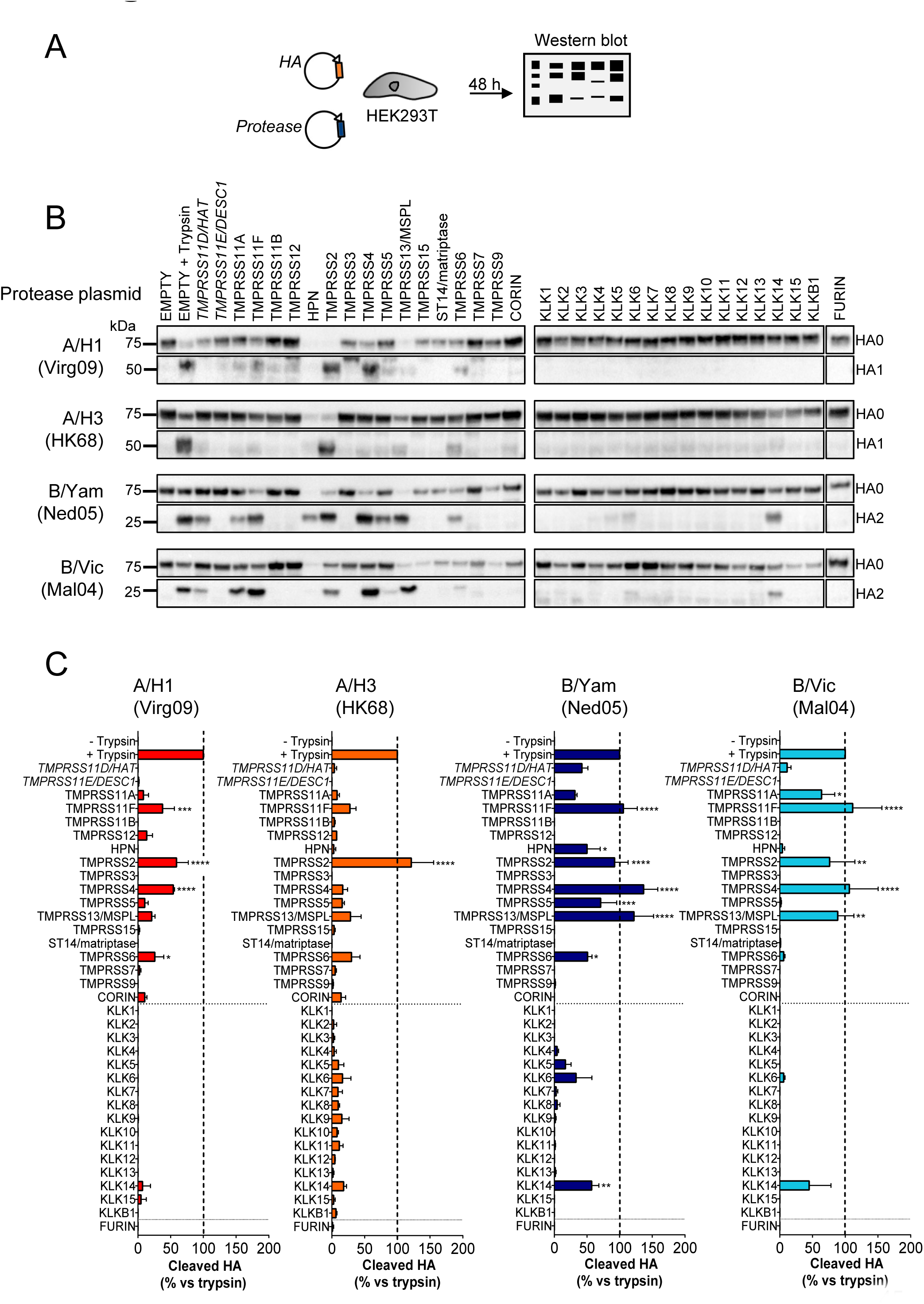
IBV HA0 is efficiently cleaved by a broad range of TTSPs. (A) Experiment set-up. HEK293T cells were transfected with two plasmids, one encoding IAV or IBV HA and one encoding the indicated TTSP, KLK or furin. The HA cleavage state was determined at 48 h post transfection. (B) Representative western blots showing the bands of uncleaved HA0 and cleaved HA1 or HA2. The trypsin controls (second lanes on each row), consisted of cells transfected with HA and a protease-lacking empty plasmid, and exposed to trypsin for 15 min just before cell extraction. (C) Quantitative data for TTSP- and KLK-mediated HA0 cleavage. For each HA, the intensity of the HA1 or HA2 band was normalized to clathrin and the % cleaved HA (mean ± SEM; N=3) was expressed relative to the trypsin control. *P*-value *versus* no trypsin: *≤ 0.05; **≤ 0.01; ***≤ 0.001; ****≤ 0.0001 (ordinary one-way ANOVA, followed by Dunnett’s test).

TMPRSS2 was the only protease that efficiently cleaved all four IAV/IBV HA0 proteins tested. It was the only protease with high activity on A/H3 HA0 (Fig. 2B and 2C), while A/H1 HA0 was equally well cleaved by TMPRSS4. On the other hand, the two IBV HA0 proteins were efficiently processed by four TTSPs (Fig. 2B and 2C), namely TMPRSS11F, TMPRSS2, TMPRSS4, and TMPRSS13/MSPL. Besides, IBV HA0 was cleaved by TMPRSS11A and TMPRSS11D/HAT, yet low expression of the latter protease (see above) precluded conclusions on cleavage efficiency. B/Yam HA0 was also recognized by TMPRSS5 and TMPRSS6. One TTSP, namely Hepsin, generated weak bands for both HA0 and HA1/HA2 (Fig. 2B), indicating HA degradation as observed before (42). Finally, none of the kallikreins was found to be a broad HA activator. KLK14 cleaved both IBV proteins, and KLK5 and KLK6 cleaved B/Yam HA0 with low efficacy. In summary, we demonstrated that, compared to its IAV counterpart, IBV HA0 can use a broader panel of TTSPs for efficient proteolytic cleavage.

### All HA0-cleaving TTSPs generate membrane fusion-competent IBV HA

We next examined whether HA0 cleavage translates into HA activation. We first employed a polykaryon assay (Fig. 3A) in which HeLa cells that co-express HA and protease, are exposed to acidic pH to trigger HA-mediated cell-cell fusion. IBV HA generated abundant polykaryons (Fig. 3B) when co-expressed with TMPRSS11F, TMPRSS2, TMPRSS4, and TMPRSS13/MSPL, consistent with efficient cleavage by these four TTSPs in the western blot assay (see above). A lower number of polykaryons was seen for TMPRSS11D/HAT, TMPRSS11A, Hepsin, TMPRSS5 and TMPRSS6. All these TTSPs thus generate a fusion-competent HA1-HA2 protein, meaning that they cleave IBV HA0 at the correct position. No polykaryons were formed in cells expressing KLK5, KLK6 or KLK14 (Fig. 3B). For KLK5 and KLK6, the levels of cleaved HA (Fig. 2) were probably too low to induce membrane fusion. KLK14 likely cleaves the IBV HA0 protein at an incorrect position, since the HA2 product generated by KLK14 migrated slightly slower (Fig. 2B) than that produced by trypsin or an activating TTSP.

**Figure 3.**
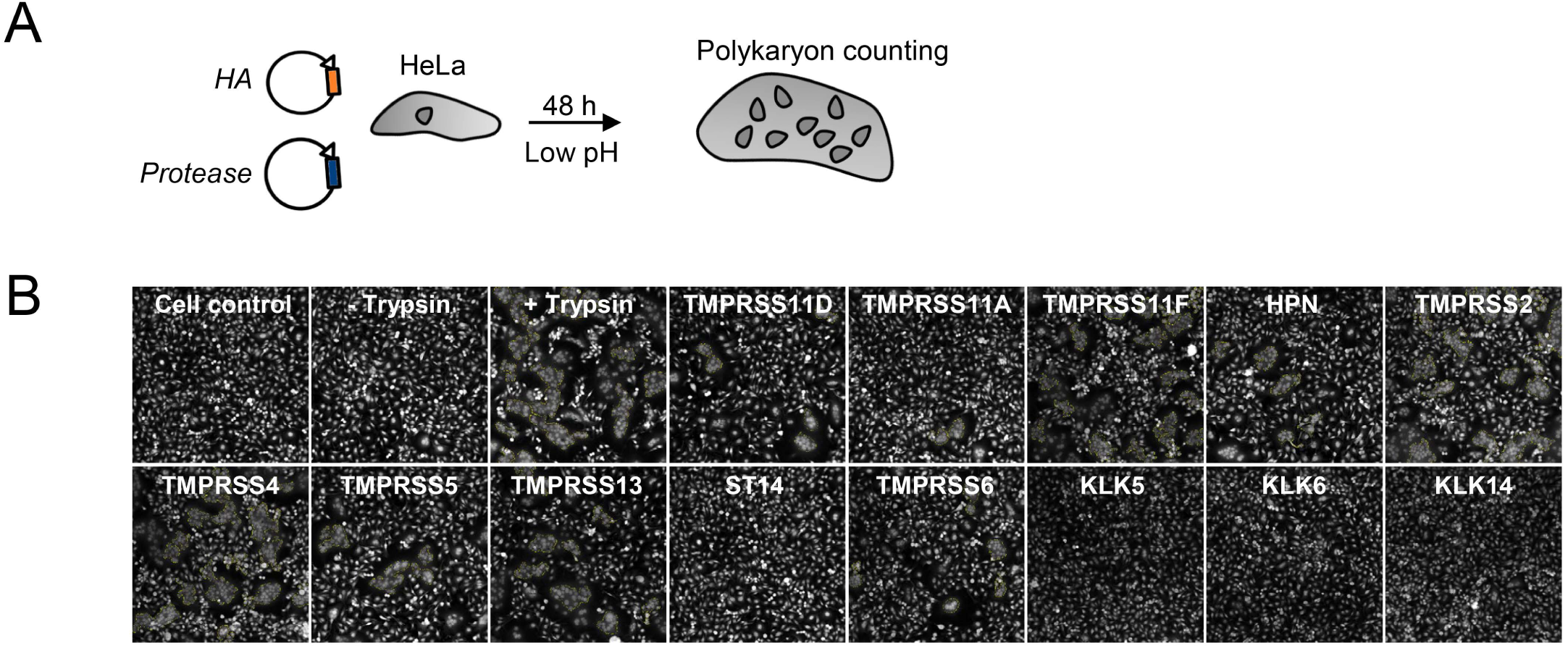
TTSP cleavage generates fusion-competent IBV HA. (A) Experiment set-up. HeLa cells expressing IBV (B/Yam) HA plus a TTSP or KLK protease were exposed to an acidic buffer 0.1 units below the fusion pH. (B) Representative photographs showing polykaryon formation in cells expressing IBV (B/Yam) HA. Polykaryon formation was similar in cells expressing B/Vic HA (not shown). The yellow contours show the polykaryons identified by high-content imaging software. The trypsin control (third photograph) received an empty (i.e. protease-lacking) plasmid and underwent HA activation by 15 min exposure to trypsin just before the acidic pulse.

### IBV shows superior TTSP activation for viral entry

To verify that the TTSPs activate HA for membrane fusion during viral entry, we employed a retroviral pseudotyping system (Fig. 4A). Each HA was combined with its cognate neuraminidase (NA) to assure efficient particle release (43, 44). Pseudoparticles were produced in the presence or absence of TTSPs, treated with PBS or trypsin and then examined for their ability to transduce target cells (Fig. 4A). In Fig. 4B and 4C, the colored bars show transduction efficiency of TTSP-activated particles, whereas the white bars show total particle infectivity upon additional treatment with trypsin.

**Figure 4.**
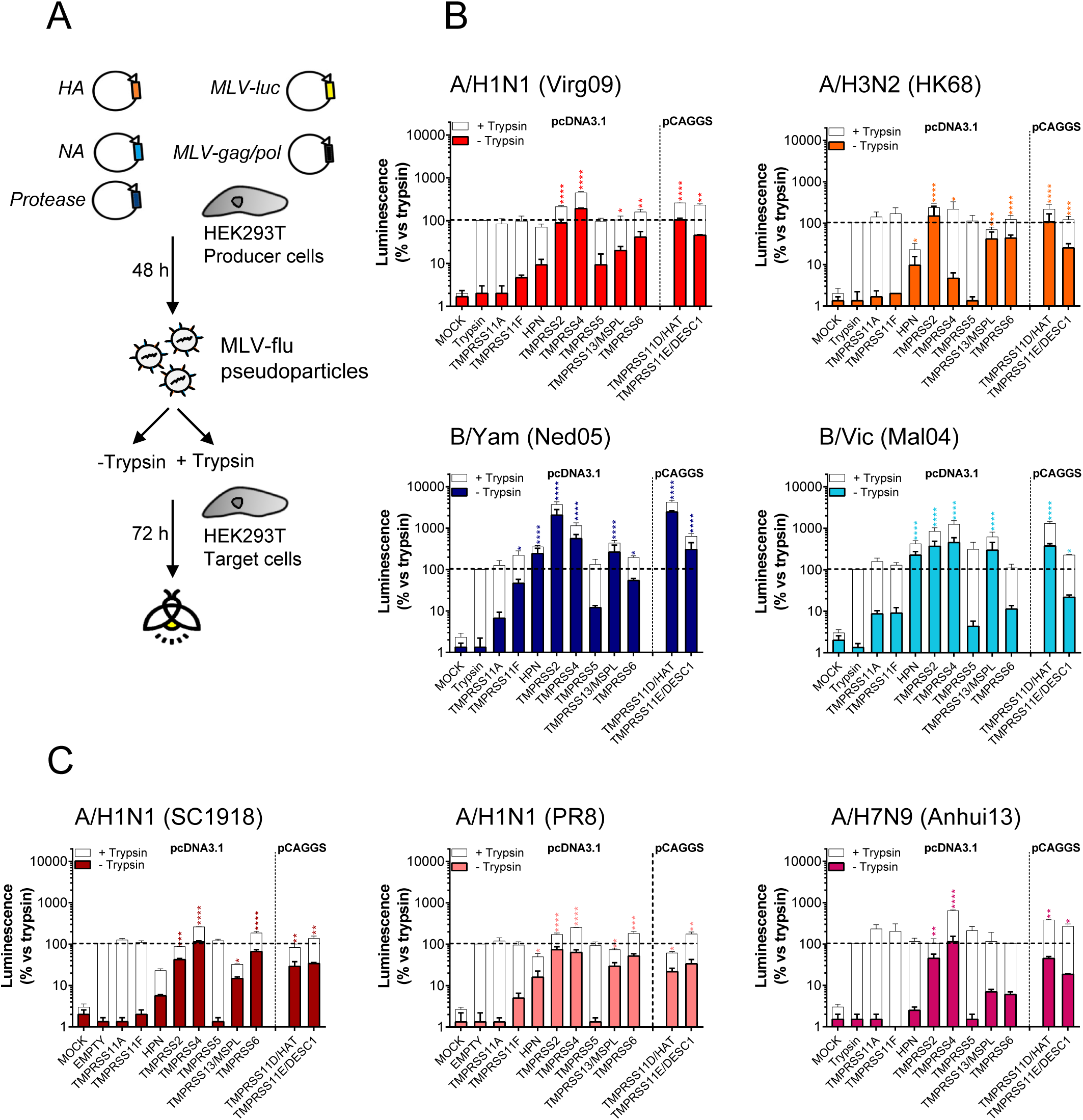
Pseudoparticles carrying IBV HA are efficiently activated by different TTSPs. (A) Experiment set-up. HEK293T producer cells were transfected with plasmids encoding the HA and NA from IAV or IBV, MLV-backbone, MLV-luciferase reporter and a TTSP enzyme. The produced pseudoparticles were left untreated (to assess TTSP-induced infectivity), or were secondarily treated with trypsin (to measure total particle infectivity). After transducing HEK293T target cells, luminescence was measured at day 3 day p.i. (B, C) Transduction efficiency (relative to the trypsin control) of TTSP-activated pseudoparticles tested as such (in color) or after additional trypsin treatment (in white). Data are the mean ± SEM (N=3 with triplicate read-outs). *P*-value *versus* no trypsin: *≤ 0.05; ***≤ 0.001; ****≤ 0.0001 (Kruskall-Wallis, followed by Dunnett’s test).

For the two seasonal IAVs, pseudoparticles produced in the presence of TMPRSS2 and TMPRSS11D/HAT (now generated from a pCAGGS-based plasmid to bypass low expression; see above) transduced cells with the same efficiency as trypsin-treated control particles (indicated with the dashed line in Figs 4B and 4C). Particles carrying A/H1 HA, but not A/H3 HA, were also fully activated by TMPRSS4. Accordingly, transduction efficiency by these particles was only slightly increased when they were treated with trypsin. In sharp contrast, IBV pseudotypes produced in the presence of Hepsin, TMPRSS2, TMPRSS4, TMPRSS13/MSPL, TMPRSS11D/HAT and TMPRSS11E/DESC1 (for B/Yam), transduced cells with markedly higher efficiency than control particles activated by trypsin (Fig. 4B). For instance, B/Yam particles activated by TMPRSS2 or TMPRSS11D/HAT generated approximately 2000-fold higher signals than the trypsin control. This superior activation of the IBV particles was also evident from the observation that subsequent trypsin treatment gave a lower signal increase for the IBV compared to the IAV particles (see white bars in Fig. 4B).

To investigate the possibility that the efficiency of HA activation might be linked to IAV virulence or pandemic potential, we included pseudotypes derived from 1918 A/H1N1 IAV and avian A/H7N9 IAV, which both possess a monobasic HA0 cleavage site. The former virus caused the devastating 1918 pandemic, and its HA protein was identified as a virulence determinant (45, 46) although the underlying mechanism is not understood (47, 48). A high case-fatality rate in humans is also seen with avian A/H7N9 IAV, which is feared for its pandemic potential (49). The three A/H1N1 pseudotypes tested, i.e. derived from 1918 IAV, 2009 pandemic virus and the laboratory strain A/PR/8/34 (PR8), as well as the A/H7N9 subtype, shared TMPRSS2 and TMPRSS4 as the most effective HA-activators (Fig. 4C). Hence, whereas TTSP activation was clearly more efficient for IBV compared to IAV pseudotypes, such a difference was not seen for highly virulent *versus* seasonal IAVs. On the other hand, 1918 IAV-pseudoparticles showed 10- and 100-fold higher transduction efficiency than the PR8- and Virg09-pseudotypes, respectively (Fig. S2) Since the three A/H1N1 pseudotypes showed comparable incorporation of HA and MLV gag and a similar level of HA0 cleavage (Fig. S2), the difference might be related to the HA fusion pH (see below) (28). An alternative explanation may be that the 1918 IAV neuraminidase, present on these pseudoparticles, may have enhanced their entry process (50).

To summarize, these polykaryon and pseudoparticle experiments established ten TTSPs as moderate to strong HA activators. Compared to the IAV proteins, IBV HA exhibits superior activation by a broader range of TTSPs.

### IBV but not IAV employs several proteases for spread in human airway-derived Calu-3 cells

Since Calu-3 cells show a similar TTSP and KLK expression profile as healthy human lungs (see above), they are a good model to study the role of these proteases in virus activation in the human respiratory tract. For the four IAV/IBV strains, i.e. Virg09 (A/H1N1), HK68 (A/H3N2), Ned05 (B/Yam) and Mal04 (B/Vic), multicycle (i.e. zanamivir-sensitive, Table S5) replication in Calu-3 cells was the same whether trypsin was added or not (*P*-value > 0.1, Fig. S3), meaning that these cells express one or more HA0-activating proteases. The four viruses were similarly inhibited by three serine protease inhibitors (see antiviral EC50 values in Table S5; a brief description of the assay is provided below this Table) but were not blocked by a furin inhibitor, consistent with furin’s inability to process monobasic HA0 cleavage sites (41). We also observed no effect with inhibitors of lysosomal cathepsins, which activate some unrelated viruses that enter by endocytosis (51). Thus, HA activation in Calu-3 cells relies on one or more serine proteases.

After optimizing the procedure for robust siRNA-mediated gene knockdown in Calu-3 cells (Fig. S4), we performed broad knockdown for the 18 TTSPs and 16 KLKs. To assess the impact of protease knockdown on IAV or IBV replication, virus was added 48 h after siRNA transfection, and virus infection was monitored at day 3 p.i. by high-content imaging of NP immunostaining (Fig. 5A). None of the siRNAs produced unwanted cytotoxic effects, the only exception being TMPRSS6 knockdown which reduced cell viability by 40% (Fig. 5B). TMPRSS2 was the only protease for which knockdown caused significant (*P*-value = 0.0001) reduction in replication of A/H1N1 and A/H3N2 virus (Fig. 5B and 5C). In sharp contrast, IBV infection was only marginally (Fig. 5B, B/Yam: *P*-value = 0.047) or not (Fig. 5B, B/Vic: *P*-value = 0.48) reduced by TMPRSS2 knockdown. Finally, IBV was not affected by single knockdown of any TTSP or KLK, nor by combined knockdown of TMPRSS2 and the most efficacious IBV HA activators identified above (Fig. 5D). Hence, the spread of IBV appears to rely on redundant proteases, concurring with the above findings that, besides TMPRSS2, several proteases effectively activate the IBV HA protein.

**Figure 5.**
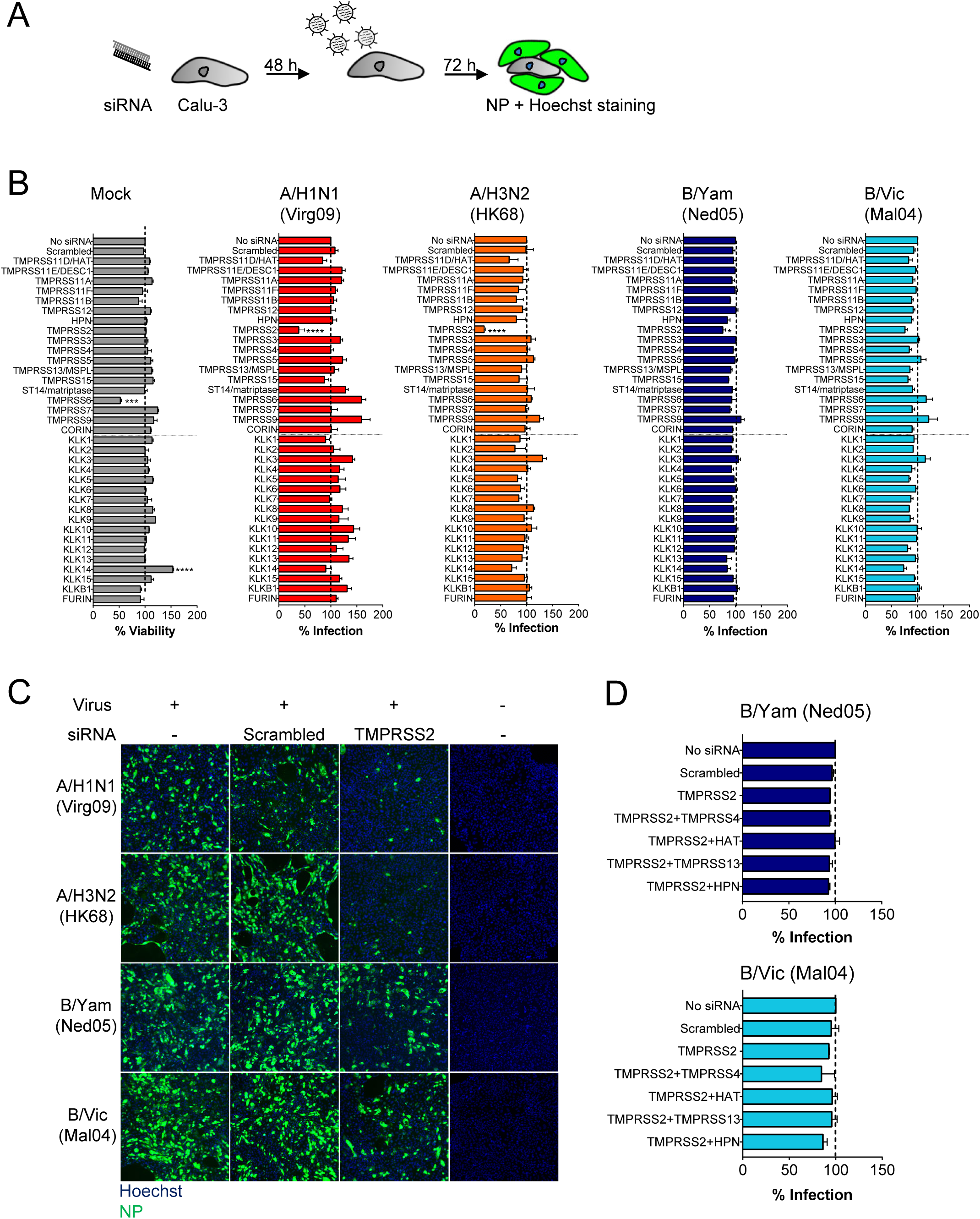
TMPRSS2 is a crucial protease for replication of IAV but not IBV. (A) Experiment set-up. Calu-3 cells were transfected with TTSP- or KLK-targeting siRNA and infected with IAV or IBV. The % infected cells was quantified by high-content imaging of NP. (B) Impact of protease knockdown on cell viability (grey bars) or virus growth (colored bars), expressed relative to the condition receiving no siRNA. Data are the mean ± SEM (N=3 with four replicates). *P*-value *versus* scrambled siRNA: *≤ 0.05; ***≤ 0.001; ****≤ 0.0001 (ordinary one-way ANOVA, followed by Dunnett’s test). (C) TMPRSS2 knockdown markedly reduces IAV but not IBV replication, as shown by NP-staining. (D) IBV infection in Calu-3 cells receiving combinations of siRNAs.

### TMPRSS4 is abundant in MDCK cells yet dispensable for spread of IBV in these cells

To assess whether some TTSPs might stand out as activators of IBV, we included Madin-Darby canine kidney (MDCK) cells which, alike Calu-3 cells, support replication of IBV in the absence of trypsin (52). This observation was also made for 1918 IAV (50, 53) which, in this regard, is unique among human IAV strains. Indeed, our seasonal IBV but not IAV viruses did not require exogenous trypsin to replicate in MDCK cells (Fig. 6A). HA0 cleavage during IBV replication was inhibited by the broad serine protease inhibitor camostat (Fig. 6B), hence executed by one or more proteases of this family. mRNA quantification in MDCK cells (see Table S3 for dog-specific primers) revealed abundant expression of *ST14/matriptase*, as observed before (54, 55) and *TMPRSS4* (Fig. 6C). Another study (50) found no *TMPRSS4* mRNA in MDCK cells, potentially related to the use of different MDCK cell lines (56). Canine TMPRSS4 was cloned into an expression plasmid to investigate its HA0-cleaving capacity. Canine TMPRSS4 proved a strong activator for IBV HA, and more effective on A/H1 HA from 1918 IAV compared to 2009 pandemic (Virg09) IAV (Fig. 6D). The lowest efficiency was seen for A/H3 HA. The polykaryon assay confirmed that fusion-competent HA was formed (pictures in Fig. 6D). The mRNA data suggested that canine TMPRSS4 or ST14/matriptase might be responsible for spread of IBV in MDCK cells. For IAV HA, published data on the role of matriptase are not consistent (55, 57, 58). Although our data thus far argued against a direct role for matriptase, an indirect role (e.g. as an activator of TMPRSS4) could not be excluded. With this is mind, we performed knockdown for canine TMPRSS4 and matriptase individually or in combination (Fig. 6E and 6F). Knockdown efficiency was >85% based on RT-qPCR for mRNA. Virus replication and HA0 cleavage in IBV-infected MDCK cells proved unaffected by knockdown of TMPRSS4, matriptase, nor a combination of the two (Fig. 6F). Thus, both in Calu-3 and MDCK cells, IBV does not seem to rely on one single protease for its replication.

**Figure 6.**
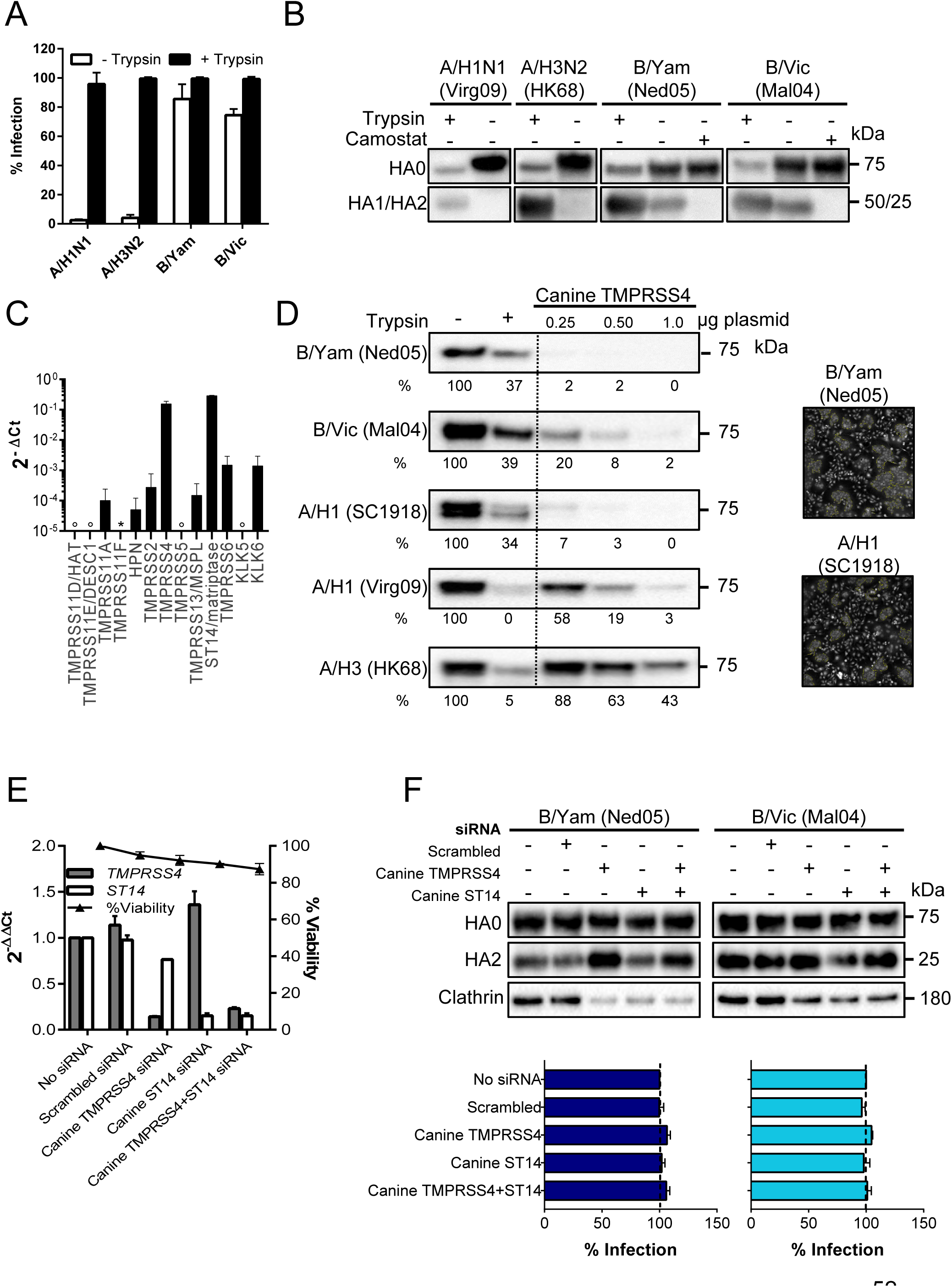
MDCK cells contain high levels of HA-activating TMPRSS4. (A, B) IAV or IBV replication in MDCK cells in the presence or absence of trypsin. Panel A: % infected cells at day 3 p.i., quantified by high-content imaging of NP. Mean ± SEM of two experiments performed in quadruplo. Panel B: HA0 cleavage state at day 3 p.i. in cells receiving 0 or 10 µM camostat. (C) Expression of TTSP and KLK proteases in MDCK cells (normalized to canine *GAPDH* and *HMBS*). °Undetermined; *no data (no dog mRNA sequence was available to design primers). (D) Canine TMPRSS4 activates the HAs of IAV and IBV. Western blot, from left to right: HA0 band in HEK293T cells receiving no protease, exogenous trypsin, or the canine TMPRSS4 plasmid at three different concentrations. Under each lane, the HA0 band intensity is given (normalized to clathrin and expressed relative to the no trypsin condition). Photographs: polykaryon formation in HeLa cells undergoing co-expression of HA and canine TMPRSS4. (E, F) Effect of TMPRSS4 and/or ST14/matriptase knockdown in MDCK cells. Panel E: mRNA levels at 24 h post siRNA transfection, normalized to canine *HMBS* and *GAPDH* and shown as the fold change *versus* untransfected control. Panel F: siRNA-transfected cells were infected with B/Yam or B/Vic virus, and at day 3 p.i., HA0 cleavage state was determined by western blot and virus infection was assessed by high-content imaging of NP.

### IBV HA exhibits a similar fusion pH as human-adapted IAV HAs

Cleavage of HA0 into HA1-HA2 activates the protein for membrane fusion, but also renders HA susceptible to inactivation at acidic pH. The human nasal cavity is mildly acidic (average pH: 6.3 in healthy adults and 5.9 in children) (30, 59). For pandemic IAV, acquirement of a more acid-stable HA is considered a prerequisite to achieve human-to-human transmissibility (27, 31, 43, 60). Since IBV is restricted to humans, we asked whether IBV possesses an acid-stable HA. Indeed, the polykaryon assay showed that the two IBV HA (i.e. B/Yam and B/Vic) proteins have a fusion pH value in the same range (5.4-5.6) as their human IAV counterparts (Fig. 7B-D), while avian A/H5 and A/H7 HAs had higher values (5.7 and 5.8, respectively; Fig. 7E), in line with previous reports (34, 43, 61, 62). On the other hand, the shape of the pH curves was distinct for IBV HA. For human-adapted IAV HAs, the curves showed a discrete pH value at which the number of polykaryons was maximal, and a steep decline at more acidic pH due to HA inactivation (63). In contrast, both IBV HAs produced high numbers of polykaryons over the entire low pH range. Hence, IBV HA is triggered for fusion at a similar acidic pH as IAV HA, however the IBV protein may be more resistant to acidic conditions. Besides, we noticed that 1918 A/H1 HA (Fig. 7D) had the highest fusion pH among the tested human IAV HAs, its value (5.7) being as high as that of avian A/H5 HA (Fig. 7E). This might explain the superior transduction efficiency of 1918 IAV pseudoparticles (see above), since an increased fusion pH enables earlier endosomal escape and more efficient infection (28).

**Figure 7.**
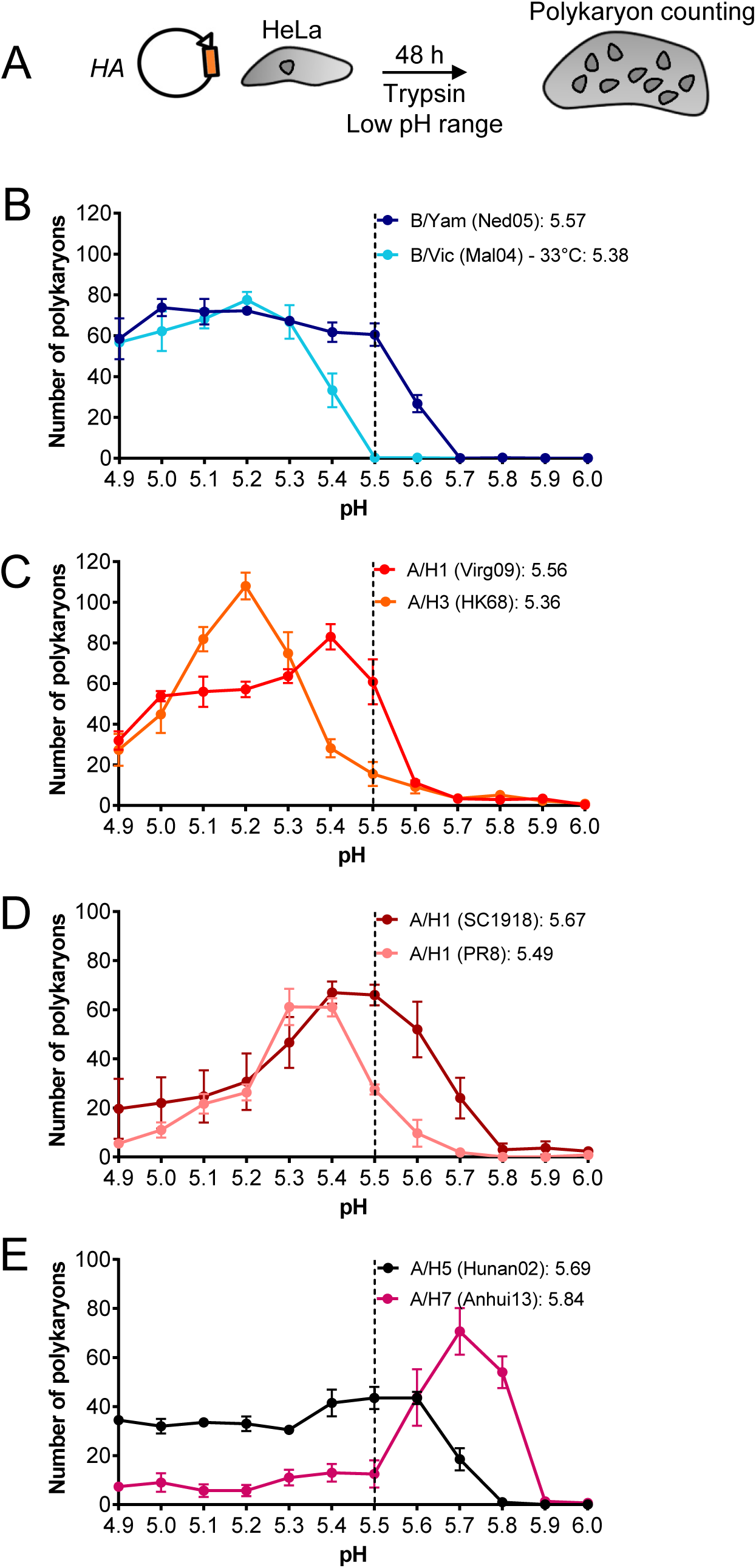
IBV HA exhibits a similar fusion pH as human-adapted IAV HAs. (A) Experiment set-up. HA-expressing HeLa cells were activated with exogenous trypsin, then exposed to a range of acidic buffers to induce cell-cell fusion. (B, C, D, E) Curves showing the number of polykaryons in function of the pH applied to trigger HA. Data are the mean ± SEM of two independent experiments, each performed in triplicate. The insets show the fusion pH values, defined as the pH at which the number of polykaryons was 50% of the maximum number.

### A temperature of 33°C is preferred by IBV HA and required for fusion by the B/Victoria lineage

The human URT has a temperature (∼30-32°C) below body temperature (35, 36) and well below that of the avian intestinal tract (∼40°C) (64, 65). Since a temperature of 33°C is preferred for propagating IBV in cell culture, we investigated whether HA has an intrinsic role in this temperature preference. Protein expression of IBV HA proved to be temperature-dependent, being highest at 33°C and gradually decreasing at higher temperatures (Fig. 8A). The 33°C preference was seen for both IBV lineages but was particularly significant for B/Vic HA (*P*-value = 0.0006 for comparison of protein levels at 33°C and 37°C). For human IAV and avian A/H5 HAs, expression was similar within the range of 33-39°C, although for A/H1 HAs, it tended to be highest at 39°C. Avian A/H7 HA manifested a clear preference for 39°C (Fig. 8A).

**Figure 8.**
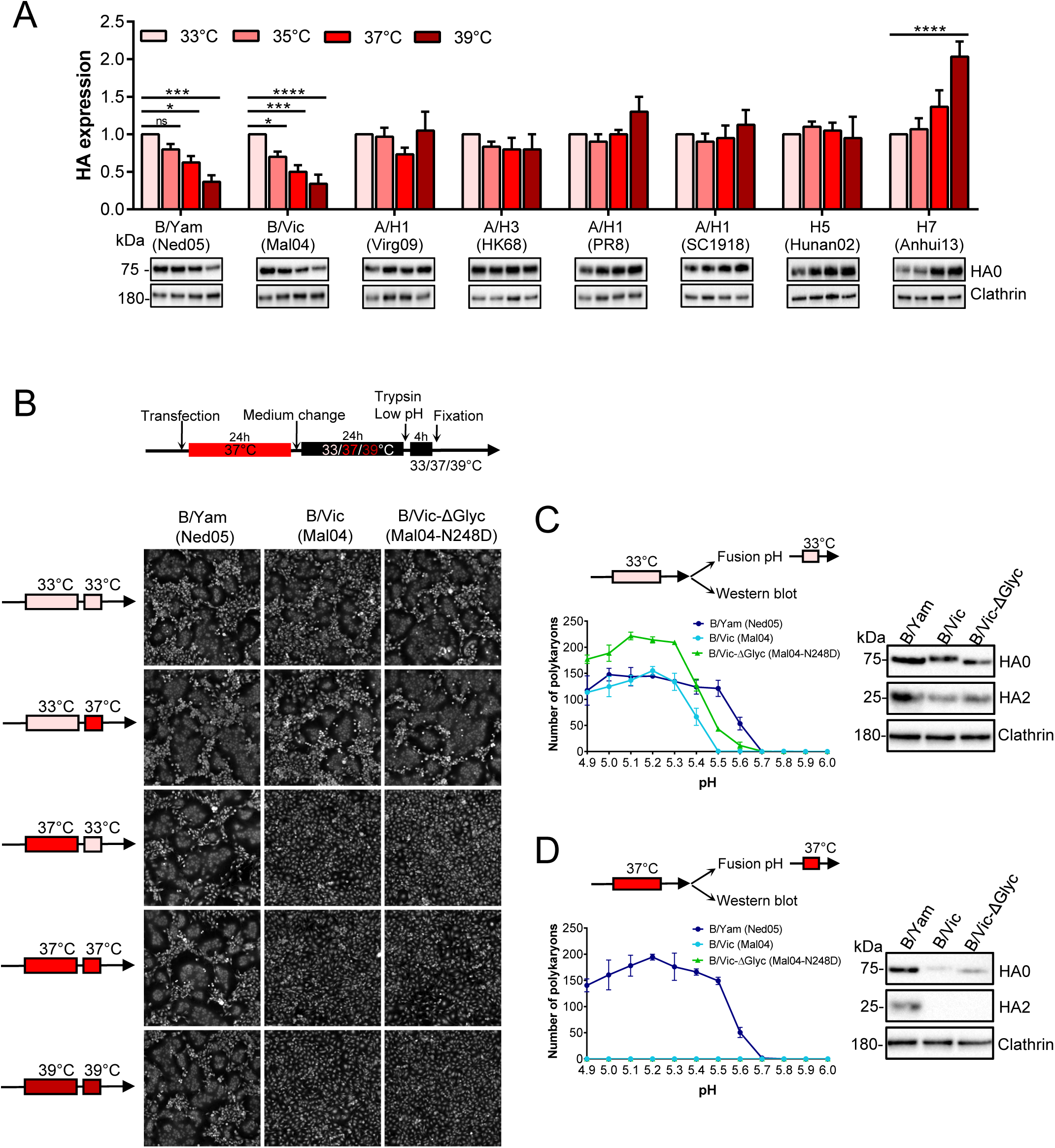
IBV HA prefers a temperature of 33°C for protein expression, explaining temperature-restricted fusion of B/Vic HA. (A) HeLa cells were transfected with the indicated HA plasmids and incubated during 48 h at four different temperatures. The graphs show quantitative western blot data for HA0 band intensity, normalized to clathrin, and expressed relative to the HA0 band seen at 33°C. Data are the mean ± SEM (N=3). *P*-values: *< 0.05, **< 0.01, ****≤ 0.0001 (ordinary one-way ANOVA, followed by Dunnett’s test). (B, C, D) Polykaryon formation in IBV HA-transfected HeLa cells exposed to different temperatures during the stages of HA expression (starting 24 h post transfection) or cell-cell fusion. Panel B: fusion induced by a pH 5.0 buffer. Panel C and panel D: number of polykaryons in function of pH, after HA expression at 33°C (C) or 37°C (D). The western blots show the corresponding HA levels after trypsin activation.

When the polykaryon assay was conducted at 37°C, B/Vic HA proved unable to induce membrane fusion at any pH tested (Fig. 8D), however abundant polykaryons were formed at 33°C (Fig. 8B and 8C). Varying the temperature during protein expression or membrane fusion (see arrow schemes in Fig. 8B) revealed that the 33°C requirement was due to inefficient HA expression at 37°C. Whether cell-cell fusion took place at 33°C or 37°C made no difference. Due to low expression at 37°C, the levels of activated HA, generated by trypsin, were undetectable (Fig. 8D). The strict 33°C requirement was not seen with B/Yam HA, since this protein generated polykaryons at all temperatures, including at 39°C (Fig. 8B). The HA of the B/Vic lineage carries a globular head glycan at residue N248, that is lacking in the B/Yam lineage (7, 66, 67). When we substituted N248 in B/Vic HA by D248, the corresponding residue in B/Yam HA, the pH threshold to induce polykaryons shifted from 5.4 to 5.5 (Fig. 8C). Polykaryons induced by the N248D-mutant were larger and more numerous compared to wild-type B/Vic HA (Fig. 8B and 8C). However, the mutation did not change the 33°C preference for protein expression and membrane fusion (Fig. 8B-D), indicating that this N248 glycan is not responsible for the temperature-sensitive phenotype of B/Vic HA.

Accordingly, pseudoparticles carrying B/Vic HA produced at 33°C had much higher infectivity compared to those generated at 37°C (*P*-value < 0.0001, Fig. 9B). Transduction efficiency was similar at both temperatures. At the cooler temperature, higher HA protein levels were visible in the producer cells and particularly in released pseudoparticles (Fig. 9C), suggesting that this reduced temperature may be required for efficient posttranslational transport and membrane incorporation of B/Vic HA.

**Figure 9.**
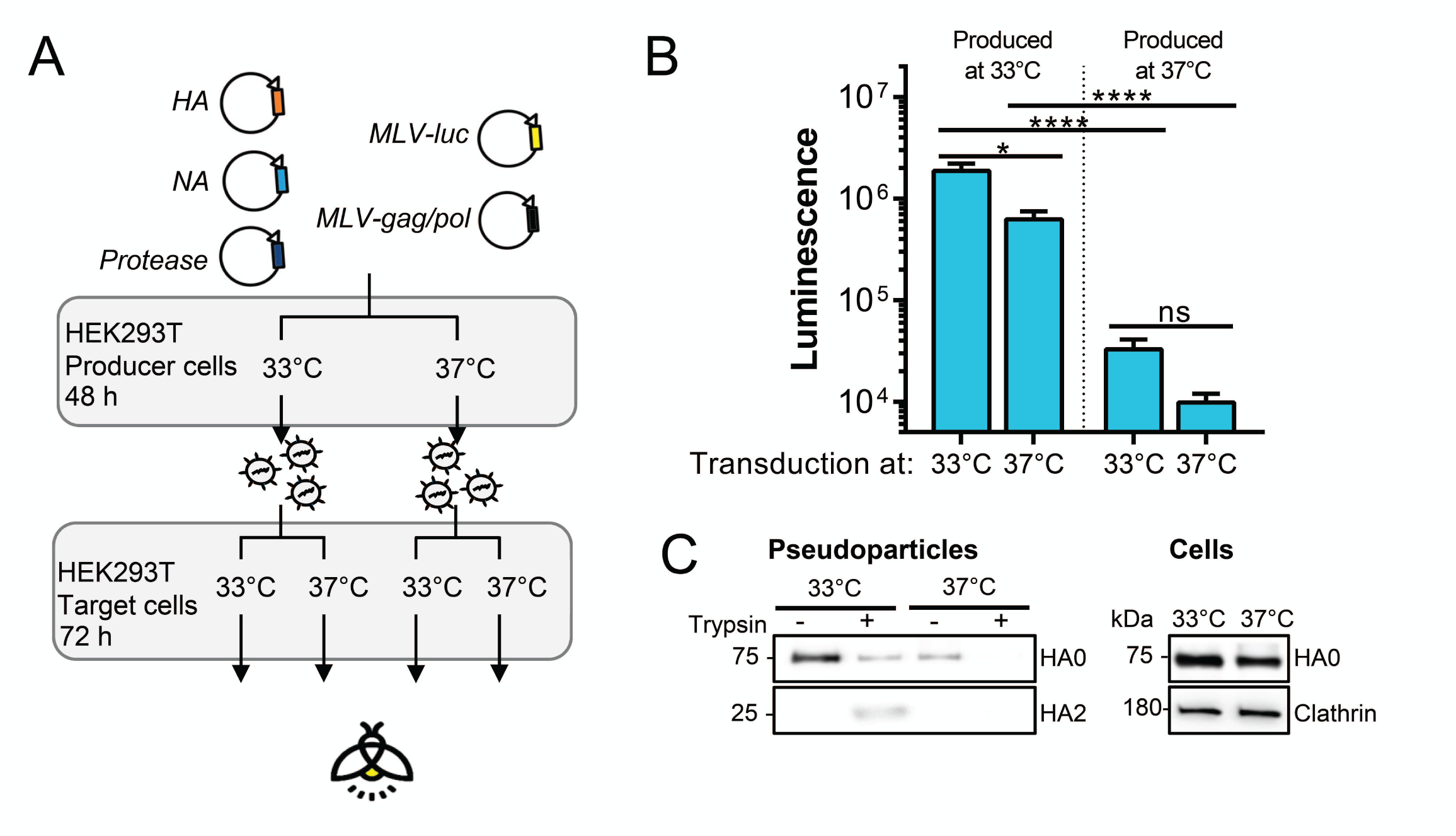
Pseudoparticles carrying B/Victoria HA require 33°C for infectivity. (A) Experiment set-up. Pseudoparticles carrying B/Vic (Mal04) HA and NA were produced at 33°C or 37°C, activated with trypsin, and transduced into HEK293T cells at 33°C or 37°C. (B) Infectivity of produced particles as measured by luminescence readout 3 days p.i. (C) HA levels in released pseudoparticles collected by ultracentrifugation and in HEK293T producer cells.

In combination, these results indicate a possible correlation between HA expression and host temperature, with IBV HA preferring 33°C, the temperature of the human URT, and avian A/H7 HA preferring the body temperature of birds. The 33°C preference was stricter for B/Vic HA than for B/Yam HA, a difference unrelated to the lineage-specific N248 head glycan.

## Discussion

Compared to the large body of literature on the HA proteins from human and zoonotic IAVs and how some of their properties reflect viral host-adaptation (8, 27), the HA of IBV is far less characterized (9). In this study, we compared the HA proteins of human IAV and IBV in terms of proteolytic activation, pH stability and temperature preference, as markers for host adaptation. Our results provide evidence for more pronounced adaptation of IBV HA to the human airways, in keeping with the long and exclusive circulation of IBV in humans.

In order to be membrane fusion-competent, HA requires cleavage by a host cell protease. Several TTSPs and KLKs have been linked to IAV HA activation (reviewed in (18)), yet a comprehensive analysis was, thus far, missing for IAV and especially for IBV. To close this gap, we compared all 18 human TTSP and 16 KLK enzymes for their capacity to cleave and activate the IAV and IBV HA proteins. Our results confirm the leading role of TMPRSS2 in activation of monobasic IAV HAs, consistent with data obtained in cell culture, knockout mice or humans (19, 20, 23). The report that TMPRSS2 proved dispensable for spread of IBV in mice (26) suggested that either TMPRSS2 is not involved at all or several redundant proteases may activate IBV HA. Our findings support the latter assumption, since the cleavability of IBV HA proved clearly superior to that of IAV. We demonstrated that both IBV HA lineages are cleaved and activated by a broad range of (in total ten) TTSP enzymes, the strongest activators being TMPRSS2, TMPRSS4, TMPRSS11F, TMPRSS13/MSPL, Hepsin and TMPRSS11D/HAT, followed by four other proteases (TMPRSS5, TMPRSS6, TMPRSS11A and TMPRSS11E/DESC1). All these TTSPs are expressed in human lung tissue and generate a functional, i.e. membrane fusion-competent, IBV HA protein. In contrast, activation of A/H1N1 and A/H3N2 HA was largely limited to TMPRSS2 and TMPRSS11D/HAT, and TMPRSS4 in the case of A/H1N1. We saw no evidence for a role of TMPRSS15/enterokinase (Hayashi 2018) or KLK5, KLK12 (68, 69) or any other kallikrein. It is possible that the KLK levels attained in our cell systems were below those applied during incubation with recombinant KLKs (68). Finally, our data do not support involvement of ST14/matriptase in cleavage of A/H1 HA (57, 58), in line with another report (55).

Limburg et al. recently showed that TMPRSS2 was crucial for IBV activation in human primary type II alveolar epithelial cells, but dispensable for spread in primary human bronchial epithelial cells and Calu-3 cells, pointing to as yet unspecified protease(s) (25). We show that single knockdown of none of the 18 TTSPs had an impact on IBV replication in Calu-3 cells, and also dual knockdown of TMPRSS2 plus another HA-activating TTSP had no effect, supporting the hypothesis that IBV can rely on redundant proteases for HA activation. This is also evident from our observation in MDCK cells, in which knockdown of the abundantly expressed TMPRSS4 nor matriptase could halt IBV replication. In contrast, replication of A/H1N1 and A/H3N2 viruses in Calu-3 cells proved strongly dependent on TMPRSS2. This is noteworthy considering that TMPRSS4 efficiently activated A/H1 HA in transfected cells and proved abundantly expressed in Calu-3 cells. How can this central role of TMPRSS2 in activation of IAV HA be explained? Unlike other TTSPs, TMPRSS2 may be present at high levels in all compartments of the constitutive secretory pathway (70) which is also followed by HA. Only TMPRSS2 appears to extensively colocalize with HA (71). Also, TMPRSS2 might not only activate HA but also the viral M2 ion channel (72). Why is IBV HA more efficiently activated than IAV HA? It is plausible that IBV HA may exhibit higher TTSP accessibility, perhaps governed by the dynamics of HA glycoprotein folding, maturation, or transport towards the cell membrane. This might be influenced by *N*-glycans in the HA head or stem (at sites very distant from the cleavage loop) having an effect on HA folding rate (73). An *N*-glycan (N8) located at the bottom of the A/H3 HA stem was shown to retard HA folding (73) and be lost when an A/H3N2 virus was passaged in TMPRSS2-knockout mice (74). Another factor is the HA0 cleavage site itself, for which two differences between IBV and IAV may be relevant: (i) at position P3 (P1 being the scissile arginine), IBV HA0 contains an extra basic lysine residue while A/H1 and A/H3 HAs carry a glutamine; and (ii) the P2’ residue is phenylalanine in IBV but leucine, a less hydrophobic residue, in human IAV HAs. Besides, the IBV HA0 cleavage loop might be structurally distinct from the A/H1 and A/H3 HA0 loops. While the latter two have been resolved (75, 76), an X-ray structure of IBV HA0 is still missing.

Since the clinical picture of IBV is similar to that of IAV (9), the high cleavability of IBV HA appears not linked to virulence. However, it assures shedding of infectious virus. Human-to-human transmissibility also requires optimal HA acid-stability, since this renders the virus more stable under mildly acidic conditions in the human URT or environment (27). For zoonotic IAVs, adaptation to the human host is associated with HA stabilization, reducing the HA fusion pH to 5.2-5.5 (31). This fits with the fusion pH values that we measured for IBV HA, i.e. 5.4 for the B/Victoria and 5.6 for the B/Yamagata lineage. Literature data for IBV are scarce but our values are in agreement with other reports (77, 78). The difference in fusion pH between the two lineages appears partially related to the B/Victoria lineage-specific N248 HA head glycan, positioned adjacent to the receptor binding site (7). Removing this *N*-glycan caused a small increase in the fusion pH threshold, pointing to an allosteric effect of the HA head domain on the fusion process, which is consistent with a role for receptor binding in fusion peptide dynamics (14).

For both IBV lineages, HA exhibits an intrinsic preference for 33°C which is particularly pronounced for the B/Victoria lineage. Analysis of additional IBV strains could validate our hypothesis that, compared to the B/Yamagata lineage, B/Victoria HA might be even better adapted to the proximal airways, given its stricter 33°C dependence and lower fusion pH. Its more stable HA could perhaps explain higher prevalence of this lineage in children (9) which typically have a more acidic nasal pH (27). A preference for cooler temperature was also reported for the hemagglutinin-esterase-fusion (HEF) protein of influenza C virus, which mainly infects humans (79). We did not observe this temperature effect for human IAV HAs, and saw the opposite pattern for avian A/H7 HA. This leads us to hypothesize that fine-tuning of HA protein expression to fit the temperature of the host target organs could be another viral adaptation strategy, besides well-known mechanisms related to receptor use, polymerase activity or immune evasion (8). The temperature-sensitive expression of IBV HA is an intrinsic feature seen in the absence of any other IBV proteins, hence it is unrelated to the reduced RNA synthesis seen with some temperature-sensitive IBV strains (80). As for the biochemical basis, for influenza C virus HEF, trimer formation and surface expression proved more efficient at 33°C than at 37°C (79). A similar temperature effect probably applies to IBV HA, since pseudoparticles carrying B/Vic HA contained much higher HA levels when produced at 33°C compared to 37°C. For IAV HA, a link exists between *N*-glycosylation and temperature sensitivity of the processes of HA folding or transport towards the cell membrane (73, 81–84). We therefore examined the role of the B/Victoria lineage-specific N248 glycan, however, removal of this HA head *N*-glycan did not change the dependence on 33°C.

Although our study was focused on the HA proteins of seasonal human IAV and IBV viruses, we also made an interesting observation for the HA of 1918 IAV. Its HA fusion pH (5.7) proved to be exceptionally high for a human IAV but identical to that of highly pathogenic avian A/H5N1 virus. This may help to explain the exceptional virulence of 1918 IAV, since IAVs with high HA fusion pH evade interferon control (85) and are more virulent in mice (86). It could also rationalize a mouse study in which the 1918 HA protein by itself generated a highly pathogenic phenotype when introduced in a contemporary backbone virus (46). Besides, we showed that TTSP activation is comparable for 1918 HA and other A/H1 HAs. Hence, the unique capacity of 1918 IAV to replicate in MDCK cells in the absence of trypsin (50, 53) seems not, or at least not entirely, explained by HA activation by TMPRSS4 or another TTSP expressed in MDCK cells. As proposed (53, 87), a role for the 1918 IAV neuraminidase is plausible.

To conclude, we demonstrated that the IBV HA protein combines broad and efficient activation capacity, favorable acid-stability and preference for the cooler temperature of the human URT. These distinct properties likely reflect host-adaptation resulting from sustained presence of this respiratory pathogen in the human population.

## Materials and methods

### Ethics statement

Lung tissue samples from eight different healthy donors were obtained under the approval of the ethical committee from the University Hospital Leuven (UZ Leuven Biobanking S51577). All patients were adult and provided written informed consent.

### Cells, media and compounds

Calu-3 (ATCC #HTB-55), A549 (ATCC #CCL-185), 16HBE [a gift from P. Hoet (Leuven, Belgium)] and HeLa (ATCC #CCL-2) cells were grown in Minimum Essential Medium (MEM) supplemented with 10% fetal calf serum (FCS), 0.1 mM non-essential amino acids, 2 mM L-glutamine and 10 mM HEPES. HEK293T cells (Thermo Fisher Scientific #HCL4517) and Madin-Darby canine kidney (MDCK) cells, a gift from M. Matrosovich (Marburg, Germany), were grown in Dulbecco’s modified Eagle’s Medium supplemented with 10% FCS, 1 mM sodium pyruvate and 0.075% sodium bicarbonate. Medium with reduced (i.e. 0.2%) FCS content was used during protease expression and virus infection experiments. Except stated otherwise, all cell incubations were done at 37°C. The following compounds were purchased from Sigma-Aldrich: N-tosyl-L-phenylalanine chloromethyl ketone (TPCK)-treated trypsin, camostat, nafamostat, aprotinin and leupeptin. Chloromethylketone, E64-d and CA-074Me were from Enzo.

### Viruses

The four human influenza virus strains and their abbreviations used in the text and figures are: A/Virginia/ATCC3/2009 (Virg09; A/H1N1 subtype; ATCC #VR-1737); A/HK/2/68 (HK68; A/H3N2 subtype) and B/Ned/537/05 (Ned05; B/Yamagata (B/Yam) lineage), both generously donated by R. Fouchier (Rotterdam, The Netherlands); and B/Malaysia/2506/04 (Mal04; B/Victoria (B/Vic) lineage; BEI resources #NR-12280). Viruses were expanded in 10-day old embryonated chicken eggs. For virus titration, a virus dilution series was added in quadruplo to Calu-3 cells. At day 3 post infection (p.i.), virus positivity was assessed by immunostaining for viral nucleoprotein (NP) (see below). Virus titers were expressed as the 50% cell culture infective dose (CCID50), calculated by the method of Reed and Muench (88).

### Plasmids

The panel of thirty-five pcDNA3.1+/C-(K)DYK plasmids containing the coding sequences for the 18 TTSPs, 16 KLKs and furin (for accession numbers and Clone IDs, see Table S1), was purchased from GenScript. The length of the ORFs was checked by PCR (Fig. S1). Expression of the different proteases was verified at 48 h post transfection of HEK293T cells, using dot blot assay. Cell lysates, prepared in RIPA buffer, were spotted onto a nitrocellulose membrane. The membrane was dried, blocked with 5% low fat milk powder in PBS, and incubated for 1 h with horseradish peroxidase (HRP)-conjugated anti-FLAG antibody (see full list of antibodies in Table S2). The dots were detected using SuperSignal™ West Femto Maximum Sensitivity Substrate (Thermo Fisher Scientific) and the ChemiDoc XRS+ System (Bio-Rad).

The expression plasmid for canine TMPRSS4 was constructed by extracting total RNA from MDCK cells, followed by reverse transcription and high-fidelity PCR using primers extended with EcoRV and XbaI sites, to allow subcloning into the pCAGEN vector provided by C. Cepko (Boston, MA) via Addgene (plasmid #11160). A similar cloning procedure was used to prepare pCAGEN vectors with the HA and neuraminidase (NA) coding sequences from Virg09, HK68, Ned05 and Mal04, starting from allantoic virus stocks. These HA proteins were >99% identical to the following GenBank sequences published on NCBI (www.ncbi.nlm.nih.gov): AGI54750.1 (for Virg09); AFG71887.1 (for HK68); AGX16237.1 (for Ned05) and CY040449.1 (for Mal04).

The coding sequences for the A/H1 HA from A/South Carolina/1/1918 (SC1918) (GenBank: AF117241.1) and the N248D-mutant form of Mal04 HA were ordered from IDT. The cDNAs for A/H5 HA (ACA47835.1) from A/duck/Hunan/795/2002 (Hunan02); and the A/H7 HA (EPI439507) and A/N9 NA (EPI439509) from A/Anhui/1/2013 (Anhui13) were purchased from Sino Biological. cDNAs were amplified by Platinum™ SuperFi™ PCR (Invitrogen) and subcloned in the pCAGEN expression vector as described above. A/H1 HA from A/PR/8/34 (PR8; Sequence ID: AYA81842.1) was subcloned into pCAGEN starting from a pVP-HA reverse genetics plasmid, kindly donated by Dr. M. Kim (Daejeon, Korea).

### Protease gene expression analysis in cells and human lung tissue

Lung tissue samples from eight different healthy donors were used. Calu-3, 16HBE, A549 and HEK293T lysates were made from three different cell passages. Total RNA was extracted using the ReliaPrep™ RNA Cell Miniprep System (Promega) and 0.5 µg RNA was converted to cDNA with the High-Capacity cDNA Reverse Transcription Kit (Thermo Fisher Scientific). BRYT Green^TM^ dye-based quantitative PCR (qPCR) was performed with the GoTaq^TM^ qPCR Master Mix (Promega) and intron-spanning primer pairs (see Table S3 for a list of all primers), in an ABI 7500 Fast Real-Time PCR system (Applied Biosciences). Expression data were normalized to the geometric mean of three housekeeping genes (*GAPDH, HMBS, ACTB*) and analyzed using the 2^-ΔCT^ method. All primers were checked for PCR efficiency and specificity by melt curve analysis. Microarray data for 108 healthy lung samples were obtained from GEO dataset GSE47460 (www.ncbi.nlm.nih.gov/geo) (89).

### Western blot assay to monitor HA0 cleavage or HA protein expression

To assess HA0 cleavage in protease-expressing cells, HEK293T cells were seeded in growth medium at 300,000 cells per well in 12-well plates, and transfected with 0.5 µg pCAGEN-HA plasmid and 0.5 µg pcDNA3.1+/C-(K)DYK-protease plasmid, using Lipofectamine 2000 (Life Technologies). Four hours later, the growth medium was replaced by medium with 0.2% FCS and cells were further incubated at 37°C or 33°C (for B/Vic HA). At 48 h post transfection, the control well was exposed to 5 µg/ml TPCK-treated trypsin for 15 min at 37°C. Cells were then lysed in RIPA buffer supplemented with protease inhibitor cocktail (both from Thermo Fisher Scientific). The lysates were boiled for 5 min in 1X XT sample buffer containing 1X XT reducing agent (both from Bio-Rad) and resolved on 4-12% Bis-Tris XT Precast 26-well gels (Bio-Rad). Proteins were transferred to polyvinylidene difluoride membranes (Bio-Rad), blocked with 5% low fat milk powder, and probed for 1 h with primary antibody followed by 45 min with HRP-conjugated secondary antibody. Clathrin served as the loading control (Table S2 for a list of all antibodies). The bands were detected and visualized as explained above for the dot blot assay. To quantify HA protein expression at different temperatures, HEK293T or HeLa cells seeded in 12-well plates were transfected with pCAGEN-HA plasmid, incubated during 48 h at 33, 35, 37 or 39 °C, then exposed to exogenous trypsin when indicated. Cell extraction and western blot analysis were carried out as above.

### Polykaryon assay to measure protease-mediated activation of HA or its fusion pH

The method was adapted from Vanderlinden *et al.* (90) to enable high-throughput format with high-content imaging. HeLa cells in black 96-well plates (20,000 cells per well) were reverse transfected with 50 ng pCAGEN-HA plasmid and 12.5 ng pcDNA3.1+/C-(K)DYK protease-plasmid using Fugene 6 (Promega). After 24 h, the growth medium was replaced by medium with 0.2% FCS and cells were further incubated at 37°C or 33°C (for B/Vic HA). Another 24 h later, the control conditions (which received HA plasmid only) were exposed for 15 min to MEM containing 5 µg/ml TPCK-treated trypsin. After gentle washing with PBS plus Ca^2+^ and Mg^2+^ (PBS-CM), the cells were exposed for 15 min to PBS-CM that had been pH-adjusted with citric acid to the required pH (i.e. a pH 0.1 unit lower than the fusion pH of that HA). After this pulse, the acidic PBS-CM was removed and growth medium with 10% FCS was added to stop the reaction. The cells were allowed to fuse during 4 h at 37°C (33°C for B/Vic HA, or another temperature if specifically mentioned), then fixated with 4% paraformaldehyde in PBS (15 min), permeabilized with 0.1% Triton X-100 (15 min) and stained for 30 min with 2 µg/ml HCS Cell Mask stain (Life Technologies). The plates were imaged using the CellInsight™ CX5 High Content Imaging Platform (Thermo Scientific). Nine images were taken per well and polykaryons were identified and counted using the SpotDetector protocol of the HCI software.

To determine the fusion pH of the different HAs, HeLa cells were transfected with the pCAGEN-HA plasmids and HA0 was cleaved with exogenous trypsin as above. During the acidic pulse, a range of acidic buffers was used having a pH between 4.9 to 6.0 with 0.1 increments. High-content imaging-based quantification of polykaryons was done as above. The fusion pH was defined as the pH at which the number of polykaryons was 50% of the maximum number (90).

### Production of protease-activated HA pseudoparticles and transduction experiments

The method to produce fLuc-expressing retroviral vectors pseudotyped with a viral glycoprotein was previously described (91). In brief, HEK293T cells were transfected in 6-well plates, using calcium phosphate precipitation, with a mixture of plasmids encoding: MLV gag-pol (1.5 µg); an MLV vector coding for fLuc (3 µg); HA- and NA-expression plasmids (both 0.75 µg); plus the protease expression plasmids specified above (0.125 µg). The two additional expression vectors pCAGGS-HAT and pCAGGS-DESC1 were described before (71). At 16 h post transfection, the medium was replaced by medium with 2% FCS. At 48 h, the pseudoparticle-containing supernatants were harvested and clarified by centrifugation. To verify that all TTSP conditions contained a similar total number of pseudoparticles, they were subsequently exposed to trypsin. Namely, one half of the supernatant was left untreated and the other half was treated with 20 µg/ml trypsin for 15 min at 37°C, after which 20 µg/ml soybean trypsin inhibitor was added. To measure particle infectivity, HEK293T target cells were seeded in 96-well plates at a density of 20,000 cells per well. One day later, the cells were exposed to 100 µl virus stock, and 6 h later, fresh medium was added. To measure fLuc activity at day 3 post transduction, the cells were lysed for 10 min in 50 µl cold PBS with 0.5% Triton-X. The lysates were transferred to a white, opaque-walled 96-well plate and, after adding fLuc substrate (Beetle-Juice (PJK) kit), the signal was read in a microplate reader (Plate Chameleon V; Hidex) using MicroWin2000 software (version 4.44; Mikrotek Laborsysteme GmbH).

### Virus replication following siRNA-mediated protease knockdown or exposure to trypsin

ON-TARGET*plus* siRNA SMARTpools targeting the thirty-five human proteases and a non-targeting (scrambled) control were ordered from Dharmacon (for catalogue numbers see Table S4). siRNAs targeting canine TMPRSS4 and canine ST14/matriptase were custom-synthesized by IDT (see sequences provided in Table S4). Cells suspended in a black 96-well plate (Calu-3: 35,000 cells per well, MDCK: 7,500 cells per well) were reverse transfected with 10 nM siRNA (each condition in quadruplo) using Lipofectamine RNAiMAX Transfection Reagent (Thermo Fisher Scientific). One day later, the transfection medium with 10% FCS was replaced by medium containing 0.2% FCS. Another 24 h later, the cells were infected with Virg09, HK68, Ned05 or Mal04 virus at a multiplicity of infection (MOI) of 100 CCID50. At 72 h p.i., immunostaining for viral NP was performed. The cells were fixated for 5 min in 2% paraformaldehyde, permeabilized for 10 min with 0.1% Triton X-100, and blocked for 4 h in 1% bovine serum albumin (BSA). Next, the plates were stained overnight at 4°C with anti-NP antibody diluted in 1% BSA (for IAV: Hytest #3IN5 at 1/2000 and for IBV: Hytest #RIF17 at 1/2000). After washing in PBS containing 0.01% Tween, Alexa Fluor^TM^ 488 secondary antibody (Invitrogen #A21424 at 1/500) was applied for 1 h at room temperature. Cell nuclei were stained with Hoechst (Thermo Fisher Scientific). The plates were imaged using the high-content platform specified above. Nine images per well were analyzed to determine the total (Hoechst) and infected (NP) cell numbers.

To measure their potential cytotoxic effects, the siRNAs were added to a parallel plate containing mock-infected cells. After five days incubation, cell viability was measured by the colorimetric MTS assay (CellTiter 96 A_Queous_ MTS Reagent from Promega) (92).

To quantify virus replication in the absence or presence of trypsin, the Calu-3 or MDCK cells were infected with virus as above, using infection medium with or without 2 µg/ml TPCK-treated trypsin, and 0.2% FCS. After 3 days incubation, the infection rate was determined by NP-staining and high-content imaging.

### Data analysis

Unless stated otherwise, data shown are the mean ± SEM of three independent experiments. Graphpad Prism software (version 7.0) was used to analyze the data and construct the graphs. One-way ANOVA or Kruskal-Wallis with post hoc test for multiple comparisons was performed to compare groups as indicated in the figure legends. To compare two groups, the unpaired Student’s t-test was used. *P*-values in the text and graphs are shown as: *≤ 0.05; **≤ 0.01, ***≤ 0.001; ****≤ 0.0001.

## Acknowledgements

Part of this research work was performed using the ‘Caps-It’ research infrastructure (project ZW13-02) that was financially supported by the Hercules Foundation (FWO) and Rega Foundation, KU Leuven. The authors wish to thank John McDonough for assisting the GEO expression analysis; Dirk Daelemans for providing the high-content imaging infrastructure; and Wim van Dam for technical assistance.

