## Supplement for "Evidence for influenza B virus hemagglutinin adaptation to the human host: high cleavability, acid-stability and preference for cool temperature"

### Table S1. Overview of the pcDNA3.1+/C-(K)DYK expression vectors used.

| | | N° | Protease | Other names | Accession number | Genscript clone ID | Length ORF (bp) |
|---|---|---|---|---|---|---|---|
| | | **1** | FURIN | | NM_002569.3 | OHu16791 | 2379 |
| Type II Transmembrane Serine Proteases | HAT/DESC subfamily | **2** | TMPRSS11D | HAT | NM_004262.2 | OHu04628 | 1251 |
| | | **3** | TMPRSS11E | DESC1 | NM_014058.3 | OHu07097 | 1266 |
| | | **4** | TMPRSS11A | HAT-like 1 | NM_182606.3 | OHu26156 | 1260 |
| | | **5** | TMPRSS11F | HAT-like 4 | NM_207407.2 | OHu31502 | 1311 |
| | | **6** | TMPRSS11B | HAT-like 5 | NM_182502.3 | OHu09192 | 1245 |
| | Hepsin/ TMPRSS2 subfamily | **7** | TMPRSS12 | | NM_182559.2 | OHu29209 | 1041 |
| | | **8** | HPN | Hepsin | NM_182983.2 | OHu08465 | 1251 |
| | | **9** | TMPRSS2 | Epitheliasin | NM_005656.3 | OHu13675 | 1473 |
| | | **10** | TMPRSS3 | | NM_032405.1 | OHu25754 | 1029 |
| | | **11** | TMPRSS4 | | NM_019894.3 | OHu19330 | 1308 |
| | | **12** | TMPRSS5 | Spinesin | NM_030770.3 | OHu04497 | 1368 |
| | | **13** | TMPRSS13 | MSPL | NM_001077263.2 | OHu21343 | 1698 |
| | | **14** | TMPRSS15 | Enteropeptidase | NM_002772.2 | OHu22654 | 3054 |
| | Matriptase subfamily | **15** | ST14 | Matriptase | NM_021978.3 | OHu19145 | 2562 |
| | | **16** | TMPRSS6 | Matriptase-2 | NM_153609.3 | OHu23829 | 2430 |
| | | **17** | TMPRSS7 | Matriptase-3 | NM_001042575.2 | OHu08737 | 2148 |
| | | **18** | TMPRSS9 | Polyserase-1 | NM_182973.2 | OHu02342 | 3174 |
| | | **19** | CORIN | TMPRSS10 | NM_006587.3 | OHu20561 | 3123 |
| Kallikreins | | **20** | KLK1 | | NM_002257.3 | OHu21121 | 783 |
| | | **21** | KLK2 | | NM_005551.4 | OHu23787 | 780 |
| | | **22** | KLK3 | | NM_001030048.1 | OHu23926 | 651 |
| | | **23** | KLK4 | | NM_004917.4 | OHu63010 | 759 |
| | | **24** | KLK5 | | NM_012427.4 | OHu25819 | 876 |
| | | **25** | KLK6 | | NM_002774.3 | OHu04090 | 729 |
| | | **26** | KLK7 | | NM_139277.2 | OHu08072 | 756 |
| | | **27** | KLK8 | | NM_144505.2 | OHu06314 | 912 |
| | | **28** | KLK9 | | NM_012315.1 | OHu19324 | 747 |
| | | **29** | KLK10 | | NM_002776.4 | OHu14898 | 825 |
| | | **30** | KLK11 | | NM_006853.2 | OHu05786 | 747 |
| | | **31** | KLK12 | | NM_019598 | OHu30141 | 762 |
| | | **32** | KLK13 | | NM_015596.2 | OHu07057 | 828 |
| | | **33** | KLK14 | | NM_001311182.1 | OHu25386 | 801 |
| | | **34** | KLK15 | | NM_001277081.1 | OHu20460 | 762 |
| | | **35** | KLKB1 | | NM_000892.4 | OHu18590 | 1911 |

**Table S2. Antibodies for western blot detection of IAV or IBV HA and dot blot detection of FLAG-tagged proteases.**

| Antibody | Company and catalog number | Host and clonality | Dilution |
|---|---|---|---|
| Anti-HA H1 (Swine flu 2009) | Sino Biological 11055-RM05 | Rabbit monoclonal | 1/4000 |
| Anti-HA H1 (A/PR/8/34) | Sino Biological 11684-RP01 | Rabbit polyclonal | 1/4000 |
| Anti-HA H1 (A/South Carolina/1/1918) | BEI resources NR-13453 | Mouse monoclonal | 1/1000 |
| Anti-HA H3 (A/Aichi/2/68) | Sino Biological 11707-T38 | Rabbit polyclonal | 1/4000 |
| Anti-HA H7 (A/Shanghai/1/2013) | Sino Biological 40104-RP02 | Rabbit polyclonal | 1/8000 |
| Anti-HA influenza B (B/Florida/4/2006) | Sino Biological 11053-R004 | Rabbit monoclonal | 1/8000 |
| Anti-clathrin heavy chain | BD Biosciences 610499 | Mouse monoclonal | 1/1000 |
| Anti-FLAG-HRP | Sigma-Aldrich A8592 | Mouse monoclonal | 1/2000 |

**Table S3. Primers for RT-qPCR quantification of human and canine *TTSP* or *KLK* mRNA in cells and lung tissue.**

| ***Protease*** | **Forward primer (5'-3')** | **Reverse primer (5'-3')** |
|---|---|---|
| *Human TMPRSS11D* | ATGGAGCATCAATGAAAAGC | GATGTTATCTCAGTTGAAGGG |
| *Human TMPRSS11E* | AACTATATGCTGAGTTTGGC | TCACCATTGATTCAAGTCTC |
| *Human TMPRSS11A* | AAGTTGTGGTAAACGAGTTG | CATGTGTTACTAATCAAGGTGG |
| *Human TMPRSS11F* | TTTGGGACTCAGTACGGCTA | GAGGCAAGGTAATAGAAAGACTTATCA |
| *Human TMPRSS11B* | CCAACAGTATCATAACTGGC | TTTCTTAGCAAAGCAGTGAG |
| *Human TMPRSS12* | CTAGCGATCCTTTAATGTGG | TCTTCTTGGTATGAGGATAGC |
| *Human HPN* | CATTGTGGCTGTTCTCCTCA | GTCCCTTCCGTCTTGTCAAA |
| *Human TMPRSS2* | GCGGATCCACCAGCTTTAT | GCAGGCTATACAGCGTAAAGAA |
| *Human TMPRSS3* | TAAGTCCTGTTGCACCAGATG | GACCAATGGCCAGTGCTAATA |
| *Human TMPRSS4* | CTTTACGAAGCAGAATGGAG | ATCATCTTCTCGGTGACTTC |
| *Human TMPRSS5* | TCAGAATAAACAGCGAAGAC | GTTGAGTTTGATGTCAGTGAG |
| *Human TMPRSS15* | GTAGTGCTCTGTGCTGGATTA | TAAATGTCGCTCTGGCTTCA |
| *Human TMPRSS13* | TCATCAACAGCAATTACACC | GATGTCTTGTCATCTGTCTC |
| *Human ST14* | CAGTCAACAACGTCAAGAAG | ATTTGTGATCCTCATGTAGC |
| *Human TMPRSS6* | TGAGGACTCCAAGAGAAAAG | CCTAGGAAATACCAGAGTAGC |
| *Human TMPRSS7* | CAGTGTTGGTCAAAGACATC | GTCTGAAATTTCCAGGTACAC |
| *Human TMPRSS9* | AAATACGCAGGAAGGAAATG | GTGAGGGAGAAGTTGTAGAG |
| *Human CORIN* | AGAATCTGTTTCACTGTCAC | ATCACAGTTTTGCTCATCAC |
| *Human KLK1* | ATTGCATCAGCGACAATTAC | CACTGACATGAACAAACTGG |
| *Human KLK2* | TTGCCTAAAGAAGAATAGCC | GAGTCTTCATCTGGTCTAAGG |
| *Human KLK3* | TTGAACCAGAGGAGTTCTTGAC | ATGAACTTGGTCACCTTCTGAG |
| *Human KLK4* | CTCATCAAGTTGGACGAATC | TCATAGAGCTTACTGCAGAC |
| *Human KLK5* | AATTCGTCCCACTAAAGATG | CACCTTTTCTGACTTAGCAC |
| *Human KLK6* | CAGCAGATGGTGATTTCC | AATCCTTCCCGTACTTCTC |
| *Human KLK7* | GCAAGATGAATGAGTACACC | GCATGAGGTCATTAACATGG |
| *Human KLK8* | CGTGGATGTTCCTGCTCTT | GGCAAGTTCTCCGCATACA |
| *Human KLK9* | CGACCCTCATCAGTGACC | GACCCTCCCATTTCCAGAG |
| *Human KLK10* | AACAACATGATATGTGCTGG | GAGCGTATGACTTTATTGATCC |
| *Human KLK11* | CCTCTACGAACATTCTTTGG | AATTTGAATCCAGGTCTCAC |
| *Human KLK12* | TCCTCACAGCGGCTCAC | GGATCTGCTCGGTCCAGTC |
| *Human KLK13* | CAAGGTTCTCAACACCAATG | TTGCCTAGGTAAACTTTGAG |
| *Human KLK14* | AAATGTTCCTCCTGCTGACA | ACGTATGGCCACCAATTATCT |
| *Human KLK15* | CAGAATCCTGTGAGGGTG | TTTGGTATAGACACCAGGC |
| *Human KLKB1* | GAAGACTGTAAGGAAGAGAAG | CACAATCTCAAAGAGTAACCAG |
| *Human Furin* | GGAAGCATGGGTTCCTCAA | GGACCGCTTCGTCACTC |
| *Human GAPDH* | CTCAGACACCATGGGGAAG | ACGGTGCCATGGAATTTGCC |
| *Human HMBS* | CCAGCTGCAGAGAAAGTTCC | TCCAAGATGTCCTGGTCCTT |
| *Human ACTB* | TGGCACCACACCTTCTACAATG | TAGCAACGTACATGGCTGGG |
| *Canine TMPRSS2* | GGTGCACCGCAAAGACTAAG | CGAGCACTTGTCCTCCATGA |
| *Canine TMPRSS4* | GGGCAGATGGGCTATGACAG | AGACAGGGTCCACTTGGGTT |
| *Canine TMPRSS11D* | CCCTTCAACTGAGATAACATCCAT | TTGCCATGGCCAATCTCCTT |
| *Canine TMPRSS11A* | AGGAACACAAGAGCCTTACCC | CAACTCGTTTACCACAACATGC |
| *Canine HPN* | CTGGCATGGAAGGGCTCTG | TAGGAAGGTCACAATGGCCC |
| *Canine TMPRSS5* | CCGGACACCACTCAACTTCT | CTGAGTTGTGGGTGTGGCT |
| *Canine TMPRSS6* | GGAGTCCAGTGCCTTCCG | GCCCCTCTCCGAAGGAGTA |
| *Canine TMPRSS13* | CACCGAGGAGAGCACAGGTAT | CTTGGACTCTGGGGCTCTCT |
| *Canine KLK5* | GGGGACATAGCAGGTACAGTG | CAGTGTCAGAGGGGTTGTCG |
| *Canine KLK6* | CTGTGGAGGGGTCCTCATTC | AGCTCTGCTCCTGGAAACTC |
| *Canine KLK13* | TTGTCAAGCGTGAGTTACCC | ACAGCATGTTGGGTGTGATCT |
| *Canine KLK14* | CACATCTAGCCCCATCGGAGT | GTCCCAGCTCCAGACAATCC |
| *Canine HMBS* | TCAGTGAGCACGTGATTGGT | ACCAGGTCGACTTCGTTCTTC |
| *Canine GAPDH* | ATTCCACGGCACAGTCAAG | TACTCAGCACCAGCATCACC |

**Table S4. Human and canine siRNAs used for protease knockdown studies in Calu-3 and MDCK cells.**

| SMARTpool Catalog Number (Dharmacon) | |
|---|---|
| *Human TMPRSS11D* | L-005894-00 |
| *Human TMPRSS11E* | L-005847-00 |
| *Human TMPRSS11A* | L-009148-00 |
| *Human TMPRSS11F* | L-032084-00 |
| *Human TMPRSS11B* | L-009116-00 |
| *Human TMPRSS12* | L-005948-01 |
| *Human HPN* | L-004332-00 |
| *Human TMPRSS2* | L-006048-00 |
| *Human TMPRSS3* | L-006049-00 |
| *Human TMPRSS4* | L-005243-00 |
| *Human TMPRSS5* | L-006051-00 |
| *Human TMPRSS15* | L-006016-00 |
| *Human TMPRSS13* | L-005973-00 |
| *Human ST14* | L-003712-00 |
| *Human TMPRSS6* | L-006052-00 |
| *Human TMPRSS7* | L-023574-00 |
| *Human TMPRSS9* | L-019463-00 |
| *Human CORIN* | L-005831-00 |
| *Human KLK1* | L-005906-00 |
| *Human KLK2* | L-005913-00 |
| *Human KLK3* | L-005914-01 |
| *Human KLK4* | L-005915-00 |
| *Human KLK5* | L-005916-00 |
| *Human KLK6* | L-005917-00 |
| *Human KLK7* | L-005918-00 |
| *Human KLK8* | L-005919-00 |
| *Human KLK9* | L-005920-00 |
| *Human KLK10* | L-005907-00 |
| *Human KLK11* | L-005908-00 |
| *Human KLK12* | L-005909-00 |
| *Human KLK13* | L-005910-00 |
| *Human KLK14* | L-005911-00 |
| *Human KLK15* | L-005912-00 |
| *Human KLKB1* | L-005921-00 |
| Non targeting control | D-001810-10 |

| Custom synthesized canine siRNA sequences (IDT) | |
|---|---|
| *Canine TMPRSS4* | UGGAUGUUGUUGGAAUCACAGAGAA |
| *Canine ST14/matriptase* | CAGAAGAUCUUCAAUGGCUACCUGA |
| Non targeting control | IDT catalog #51-01-14-03 |

**Table S5. Antiviral activity of protease inhibitors and influenza virus inhibitors in Calu-3 cells infected with IAV or IBV.**

| Compound | Antiviral activity ($EC_{50}$ in µM) | | | | Cytotoxicity ($CC_{50}$ in µM) |
|---|---|---|---|---|---|
| | A/H1N1 (Virg09) | A/H3N2 (HK68) | B/Yam (Ned05) | B/Vic (Mal04) | |
| **Camostat**<br>Broad inhibitor of serine proteases | 1.9 ± 0.2 | 1.6 ± 0.2 | 1.3 ± 0.2 | 5.0 ± 4.0 | >50 |
| **Nafamostat**<br>Broad inhibitor of serine proteases | 0.6 ± 0.2 | 0.3 ± 0.03 | 1.4 ± 0.2 | 19 ± 12 | >50 |
| **Aprotinin**<br>Broad inhibitor of serine proteases | 29 ± 3 | 17 ± 1 | 13 ± 1 | >50 | >50 |
| **Leupeptin**<br>Broad inhibitor of serine and cysteine proteases | >25 | >25 | >25 | >25 | >50 |
| **Chloromethylketone**<br>Furin inhibitor | >25 | >25 | >25 | >25 | >50 |
| **E64-d**<br>Inhibitor of cysteine proteases | >25 | >25 | >25 | >25 | >50 |
| **CA-074Me**<br>Inhibitor of cathepsin B | >25 | >25 | >25 | >25 | >50 |
| **Ribavirin** | 31 ± 2 | 9.5 ± 0.6 | 18 ± 1.0 | 18 ± 2.8 | >50 |
| **Zanamivir** | 6.8 ± 1.7 | <0.2 | 0.3 ± 0.1 | 33 ± 10 | >50 |

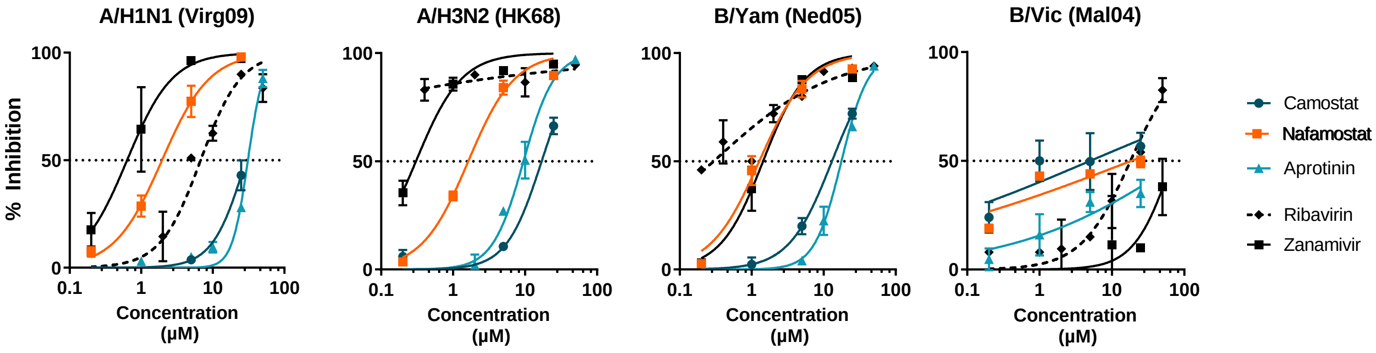


**Methodology**

Serial dilutions of the compounds were added to Calu-3 cells just before infection with IAV or IBV (MOI: 100x$CCID_{50}$). At day 3 p.i., the cells were immunostained for viral NP, after which high-content imaging was performed. Data were expressed relative to the virus control set at 100%, and plotted in GraphPad Prism using non-linear regression. $EC_{50}$ values were calculated from the obtained dose-response curves. The mean ± SEM are shown. Antiviral activity is expressed as the 50% effective concentration ($EC_{50}$), defined as the compound concentration producing 50% inhibition of virus replication. Compound cytotoxicity is expressed as the 50% cytotoxic concentration ($CC_{50}$), estimated by MTS cell viability assay on mock-infected cell cultures.

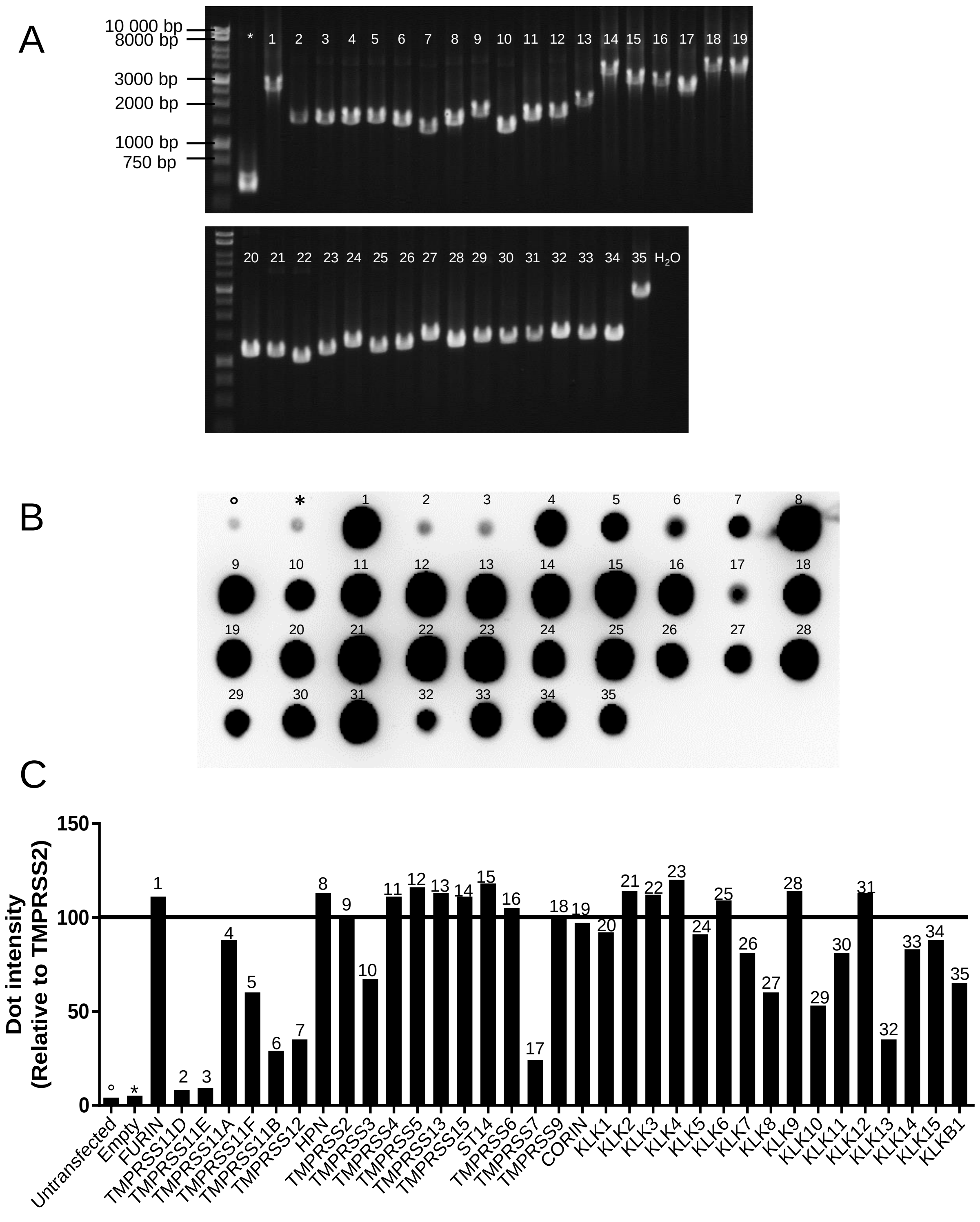


**Fig. S1. PCR control and protein expression levels for the TTSP- and KLK-expression plasmids.** (A) The correct length of the ORFs was checked by PCR using a CMV promoter-directed forward primer and flag/DYK-directed reverse primer. (B) Successful protein expression after plasmid transfection in HEK293T cells was checked by anti-flag dot blot assay. °Untransfected cell control; *cells transfected with empty vector. (C) Quantification of the dots shown in panel B; dot intensities were calculated relative to the TMPRSS2 dot, set at 100%. The numbers correspond to the plasmid numbers shown in Table S1.

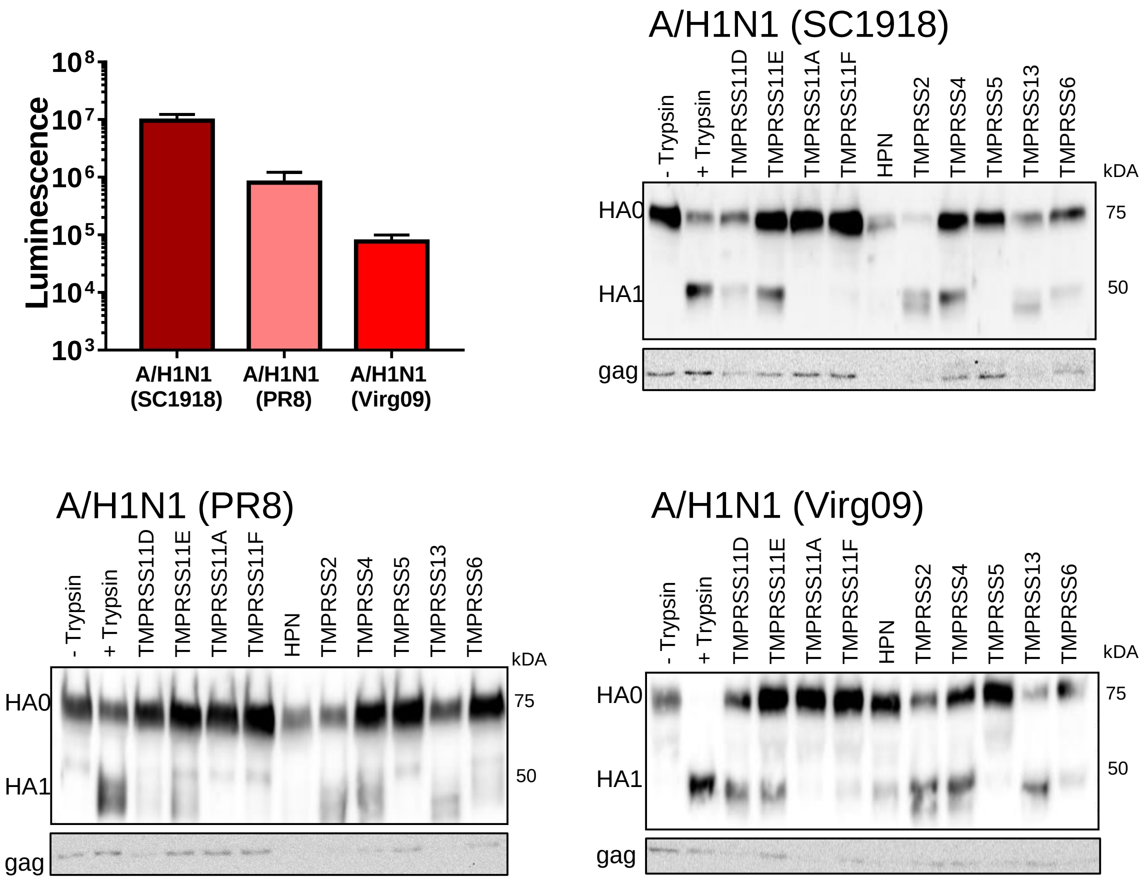


**Fig. S2. Transduction efficiency of A/H1N1 pseudoparticles and their HA cleavage state.** Pseudoparticles treated with trypsin were used to transduce HEK293T cells, luminescence was measured 3 days p.i. To assess Gag incorporation and HA cleavage status, particles were purified by ultracentrifugation through a 20% sucrose cushion. The pellets were resuspended in SDS-PAGE sample buffer and analyzed by western blot. MLV-Gag was detected using an anti-MLV-p30 mouse antibody (Acris).

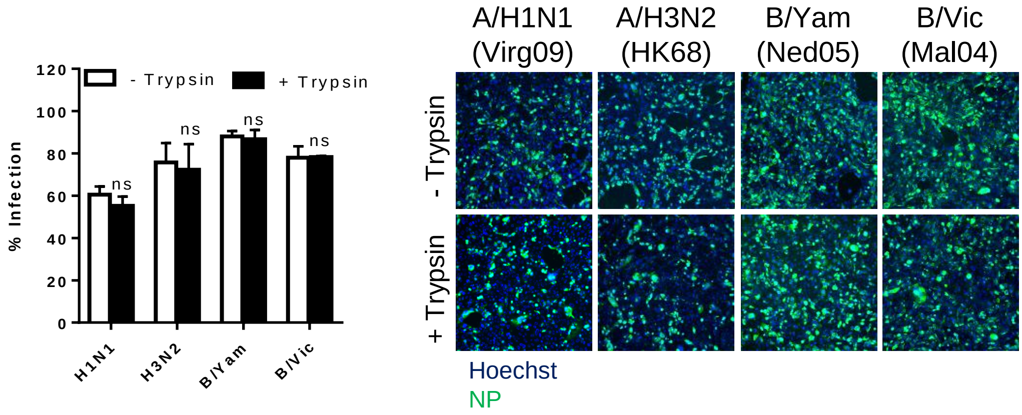


**Fig. S3. Trypsin-independent replication of IAV and IBV in Calu-3 cells.** At day 3 p.i., virus was quantified by immunostaining for viral NP followed by high content imaging. The bar graph shows the mean ± SEM ; ns: not significant (*P*-value > 0.1 by unpaired t-test comparing the condition in the presence *versus* absence of trypsin).

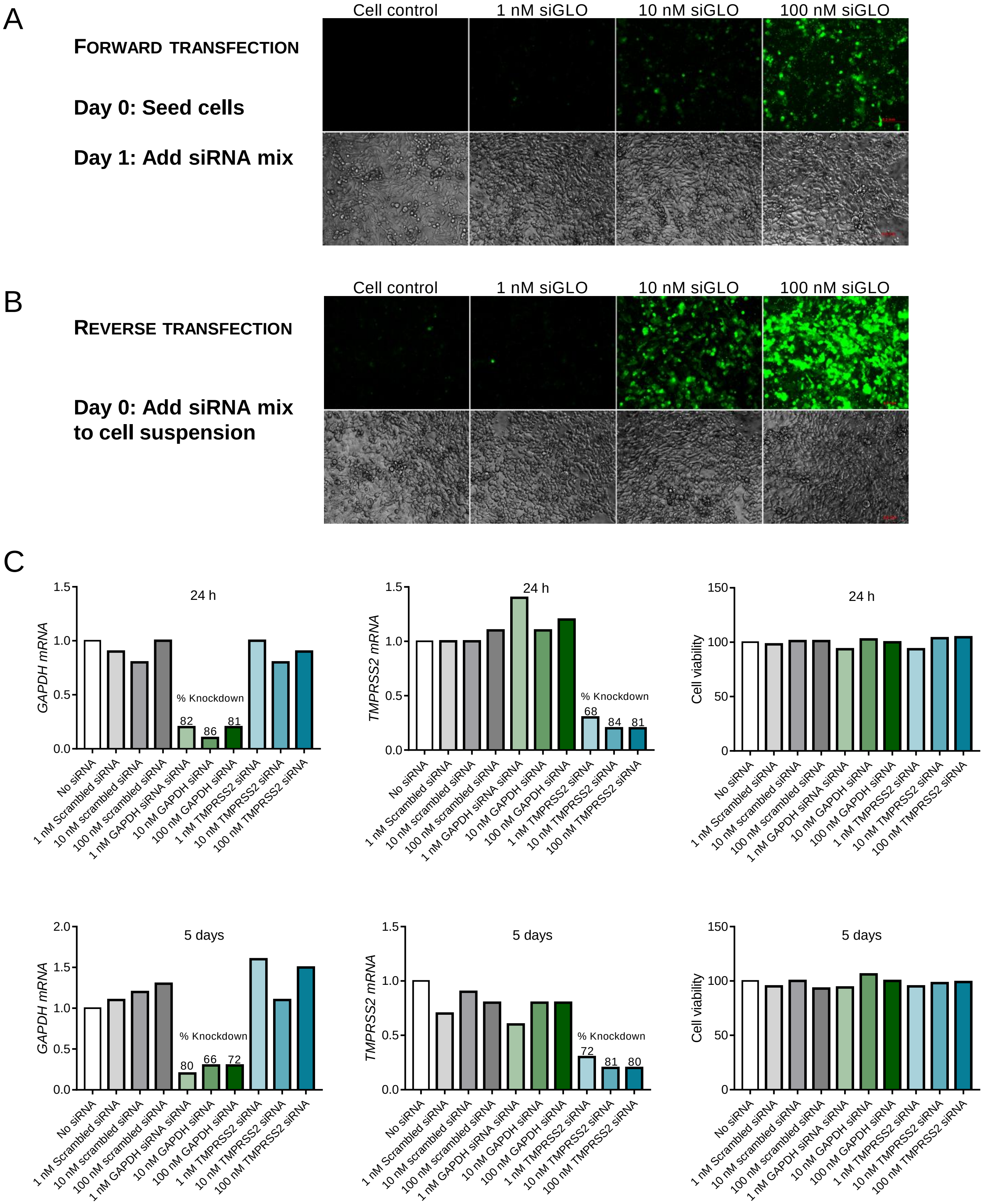


**Fig. S4. Optimization of siRNA transfection in Calu-3 cells.** Calu-3 cells were transfected with different amounts of siGLO Green Transfection Indicator (Dharmacon) using two different approaches. (A) cells were plated on day 1 and transfection complexes were added on day 2 (forward transfection). (B) cells and transfection mixes were added at the same time (reverse transfection). Cells were imaged by fluorescence microscopy at 24 h post transfection. (C) Analysis of *GAPDH* and *TMPRSS2* mRNA knockdown at 24 h post siRNA reverse transfection. The expression levels were normalized to the housekeeping genes *HMBS* and *ACTB*. Relative differences were calculated as the fold change *versus* the untransfected control sample (no siRNA).
